# Feedback regulation of steady-state epithelial turnover and organ size

**DOI:** 10.1101/161943

**Authors:** Jackson Liang, Shruthi Balachandra, Sang Ngo, Lucy Erin O’Brien

**Affiliations:** Department of Molecular and Cellular Physiology, Stanford University School of Medicine, Stanford, CA 94305.

## Abstract

Epithelial organs undergo steady-state turnover throughout adult life, with old cells being continually replaced by the progeny of stem cell divisions^1^. To avoid hyperplasia or atrophy, organ turnover demands strict equilibration of cell production and loss^2-4^. However, the mechanistic basis of this equilibrium is unknown. Using the adult *Drosophila* intestine^5^, we find that robustly precise turnover arises through a coupling mechanism in which enterocyte apoptosis breaks feedback inhibition of stem cell divisions. Healthy enterocytes inhibit stem cell division through E-cadherin, which prevents secretion of mitogenic EGFs by repressing transcription of the EGF maturation factor *rhomboid*. Individual apoptotic enterocytes promote divisions by loss of E-cadherin, which releases cadherin-associated β-catenin/Armadillo and p120-catenin to induce *rhomboid*. Induction of *rhomboid* in the dying enterocyte triggers EGFR activation in stem cells within a discrete radius. When we block apoptosis, E-cadherin-controlled feedback suppresses divisions, and the organ retains the same number of cells. When we disrupt feedback, apoptosis and divisions are uncoupled, and the organ develops either hyperplasia or atrophy. Altogether, our work demonstrates that robust cellular balance hinges on the obligate coupling of divisions to apoptosis, which limits the proliferative potential of a stem cell to the precise time and place that a replacement cell is needed. In this manner, localized cell-cell communication gives rise to tissue-level homeostatic equilibrium and constant organ size.

## MAIN TEXT

Over an animal’s lifetime, mature organs undergo repeated rounds of cell turnover yet are able to remain the same approximate size. This remarkable ability implies the existence of robust mechanisms to ensure that turnover is zero-sum, with precisely equal rates of cell production and loss ^1,2,10-12^. In most organs, the production of new cells ultimately depends on the divisions of resident stem cells. Although much is understood about how excessive or insufficient divisions lead to disease, little is known about how equal rates of division and loss are sustained during the steady-state turnover of healthy tissues.

We investigated the regulation of turnover using the epithelium of the adult *Drosophila* midgut ^8,9^. To establish whether production of new cells equals loss of old cells, we examined the kinetics of cell addition and loss using *escargot* flp-out (*esg*^*F/O*^) >*GFP* labeling (Fig. 1a-e, Extended Data Fig. 1) ^13^. Upon 29°C temperature shift, all undifferentiated midgut cells are labeled by permanent, heritable GFP expression. Mature cells that were present before induction remain unlabeled, while mature cells that arise after induction inherit GFP expression from their progenitors. Focusing on the midgut’s R4ab region (also known as P1-2; Extended Data Fig 1b-e)^14,15^, we found that total (DAPI^+^) cells remained near-constant over time, while the number of newly added, GFP^+^ cells increased linearly (Fig 1e, Extended Data Fig 1g). At 4 days post-induction, virtually all cells in the R4ab region were GFP^+^, signifying that complete cell replacement had occurred. We conclude that production of new cells quantitatively equals loss of old cells.

**Fig. 1.**
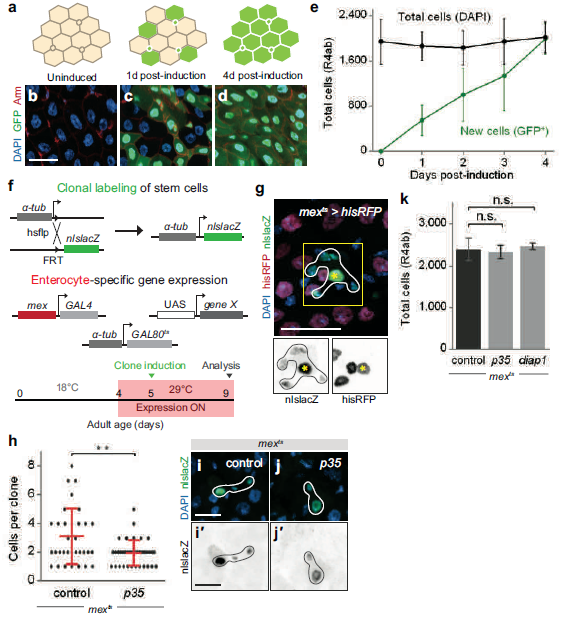
Enterocyte apoptosis regulates the rate of stem cell division for homeostatic maintenance of overall cell number. **a-e,** The midgut R4ab compartment undergoes complete cell turnover in 4 days. **a,** Cartoon of *esg*^*F/O*^*>GFP* labeling strategy to quantify kinetics of turnover. Before induction, all progenitor cells (stem and enteroblast cells, small circles) and mature enterocytes (hexagons) are unmarked (tan). Upon induction by 29°C temperature shift, all progenitor cells express GFP (green). Postinduction, newly generated enterocytes inherit GFP expression from progenitors, while preexisting enterocytes stay unmarked. Cell turnover is complete when all cells are GFP^+^. See also Extended Data Fig 1. **b-d**, Representative images of *esg*^*F/O*^*>GFP* midguts before induction, 1 day post-induction, and 4 days post-induction. DAPI (blue) marks all nuclei. Armadillo (Arm, red) marks cell boundaries. **e,** Quantification of total (DAPI^+^) and new (GFP^+^) cells in the R4ab compartment over time ^14,15^. Number of total cells stays near-constant. Number of GFP^+^ cells increases linearly. After 4 days, number of GFP^+^ cells becomes equal to total cells. Each time point represents 3 midguts. **f-g,** Tracing stem cell divisions in a background of genetically manipulated enterocytes. **f,** Clonal labeling of stem cell divisions is induced by a brief pulse of FRT recombination that reconstitutes a split *a-tub-nlslacZ* transgene. Enterocyte-specific gene expression is turned on by 29°C shift that permits *mexGAL4* to drive expression of UAS-*gene X* (*mex*^*ts*^*>gene X*). See also Extended Data Fig. 2. **g**, Validation of genetic system using *mex*^*ts*^*>his2av::RFP*. β-galactosidase marks a stem cell clone (outlined) in a background of His2av::RFP^+^ enterocytes. Within the 5-cell clone, only the enterocyte (yellow asterisk, polyploid) expresses *his2av::RFP*. **h-k,** Blocking enterocyte apoptosis causes fewer stem cell divisions but does not change total cell number. **h,** Clone size analysis in the R4ab compartment. Each point is the number of cells in one β-gal-marked clone. Data pooled from 4-5 midguts per genotype. Values are means ± S.D. Mann-Whitney test, *p*=0.009. **i-j,** Images show average-sized clones. **k,** Total R4ab cell counts are comparable for control, *mex*^*ts*^*>p35* and *mex*^*ts*^>*diap1* midguts. Cells were counted 4 days post-induction. N=4 midguts per genotype. Values are means ± S.D. Unpaired t-test, *p*>0.05. One of three representative experiments is shown in each graph. All scale bars are 25 μm.

Most cells in the midgut epithelium are polyploid enterocytes ^16^. Each enterocyte is the product of one asymmetric stem cell division; the enterocyte lineage contains no transit-amplifying cells (Extended Data Fig. 1a) ^9^. To probe the relationship between cell production and loss, we devised a system to manipulate enterocytes and perform concomitant lineage tracing of stem cells by combining enterocyte-specific *mexGAL4;GAL80*^*ts*^ (*mex*^*ts*^) ^17,18^ with the clonal labeling system split-*nlsLacZ* ^6,9,19-21^ (Fig 1f; Extended Data Fig. 2). This two-pronged system was tested by de-repressing *mex*^*ts*^*>his2av::RFP* with 29°C temperature shift and then inducing mitotic recombination and β-galactosidase expression in stem cells via a brief 38.5°C heat shock. As expected, His::RFP marked all enterocytes (polyploid cells), both within and outside of β- galactosidase-marked stem cell clones, but did not mark non-enterocyte, diploid cells (Fig 1g).

**Fig. 2.**
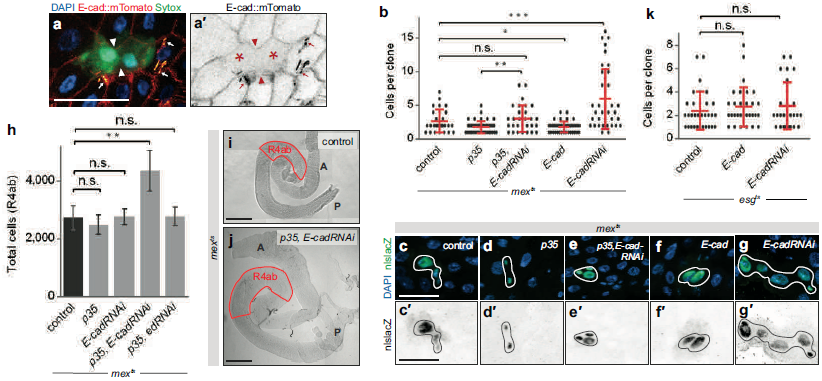
Homeostatic size control requires *E-cad* on enterocytes, but not on stem cells. **a**, Dying enterocytes lose junctional E-cad. Endogenous E-cad tagged with mTomato (red hot LUT) localizes to the lateral membranes of healthy enterocytes but not dying, Sytox^+^ enterocytes (green; asterisks in **a’**). E-cad loss is most pronounced where two dying enterocytes are juxtaposed (arrowheads). In these live images, trachea exhibit bright, yellow-orange autofluorescence (arrows). Representative images are shown from two independent experiments; N=4 midguts per experiment, analyzed 6 days post-eclosion. Scale bar, 25 μm. **b-g,** Enterocyte E-cad is necessary and sufficient to repress stem cell divisions. **b,** Clone size analysis in the R4ab compartment using schema/timing in Fig. 1f. Stem cell clones are larger in *mex*^*ts*^*>p35, E-cadRNAi* midguts compared to *mex*^*ts*^*>p35* alone (Mann-Whitney test, *p*=0.0034). Clones are also larger in *mex*^*ts*^*>E-cadRNAi* midguts compared to control (*p*=0.0004). Clones are smaller in *mex*^*ts*^*>E-cad* compared to control (*p*=0.04). **c-g,** Images show average-sized clones. Scale bars, 25 μm. **h**, Enterocyte *E-cad* is necessary to maintain constant cell number during apoptotic inhibition. Total cell number is normal in *mex*^*ts*^*>p35* and *mex*^*ts*^*>E-cadRNAi* midguts but increases by ~70% in *mex*^*ts*^*>p35, E-cadRNAi* midguts (unpaired t-test, *p*=0.007) after 4 days. This increase is unlikely caused by non-specific loss of cell-cell adhesion because total cell number remains normal in *mex*^*ts*^*>p35, echinoid(ed)RNAi* midguts. N=4 midguts per genotype; means ± S.D shown. **i-j,** Enterocyte *E-cad* is necessary to maintain constant organ size during apoptotic inhibition. Organ size is larger in *mex*^*ts*^>*p35, EcadRNAi* midguts compared to controls. A, anterior; P posterior. Scale bars, 200 μm. See also Extended Data Fig. 3a. **k,** Stem cell/enteroblast *E-cad* does not control stem cell divisions. Clone size analysis in R4ab as in Fig. 1f, except that *esgGAL4* was used to manipulate *E-cad* in stem and enteroblast cells. Stem cell clones are comparably sized in *esg*^*ts*^ control midguts, *esg*^*ts*^ *>E-cad* midguts, and *esg*^*ts*^ *>E-cadRNAi* midguts (Mann-Whitney test, *p*>0.05). In **b** and **k**, data are pooled from 4-5 midguts per genotype. Values are means ± S.D. Each graph comprises data from three representative experiments.

Using this two-pronged system, we blocked enterocyte apoptosis and assessed the impact on stem cell divisions. Expression of the potent apoptotic inhibitor *p35* in enterocytes (*mex*^ts^*>p35*) resulted in fewer stem cell divisions, as indicated by smaller clone sizes (Fig 1h-j). Slowing down stem cell divisions could be a compensatory strategy to preserve overall cell number. Indeed, total cell numbers remained constant when apoptosis was blocked by p35 or by the native caspase inhibitor Diap1 (Fig 1k) after four days. The physical dimensions of apoptosis-inhibited midguts were similar to control guts (Extended Data Fig 3a), and epithelial tissue architecture remained intact (Extended Data Fig 4a-b, d-e). These results are supported by a prior report that fewer midgut cells incorporate BrdU in animals with impaired caspase activation ^22^. Altogether, these findings imply that enterocyte apoptosis regulates the rate of stem cell divisions to homeostatically maintain constant cell number and organ size.

**Fig. 3.**
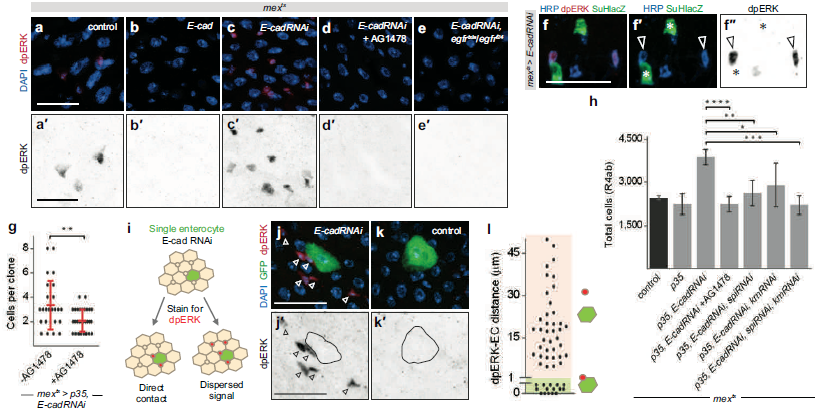
Enterocyte E-cad inhibits stem cell EGFR via a dispersed signal for homeostatic size control. **a-f,** Enterocyte E-cad inhibits stem cell EGFR signaling. **a-c,** Immunostaining for diphosphorylated ERK (dpERK). dpERK^+^ cells are sparse in control midguts, absent in *mex*^*ts*^*>E-cad* midguts, and abundant in *mex*^*ts*^*>E-cadRNAi* midguts. **d-e,** dpERK signal is eliminated following EGFR inhibition by oral administration of AG1478 or by temperature-induced inactivation of an *egfr*^*tsla*^*/egfr*^*f24*^ heteroallele. **f,** dpERK staining is limited to stem cells (HRP^+^, Su(H)lacZ^-^; arrowheads in **f’** and **f’’**) and does not mark enteroblasts (HRP^+^, Su(H)lacZ^+^; asterisks in **f’** and **f’’**), even in *mex*^*ts*^*>E-cadRNAi* midguts. Representative images are shown from two independent experiments; N=4 midguts per genotype in each experiment, analyzed after 2 days of transgene expression. See also Extended Data Fig. 3b. **g,** EGFR activation is necessary for *E-cad*-depleted enterocytes to accelerate stem cell divisions. Clone size analysis in R4ab using schema in Fig 1f. Stem cell clones in *mex*^*ts*^*>p35, E-cadRNAi* midguts are smaller when EGFR is inhibited by AG1478. Data pooled from 4-5 midguts per genotype. Mann-Whitney test, *p*=0.008. Values are means ± S.D. **h,** Organ hyperplasia requires EGFR and the EGF ligands *spitz* (*spi*) and *keren* (*krn*). AG1478 treatment restores total cell numbers of *mex*^*ts*^*>p35, E-cadRNAi* midguts to the range of control and *mex*^*ts*^*>p35* midguts (unpaired t-test: *p*<0.0001). RNAi of either *spi* or *krn* in enterocytes reduces total cell numbers, and double RNAi of *spi* and *krn* restores total cell numbers to the normal range (*p*=0.0026, 0.046, and 0.0002 respectively). N=4 midguts per genotype, analyzed 4 days post-induction. Values are means ± S.D. See also Extended Data Fig. 3a. **i-l,** EGFR activation involves a dispersed signal. Single enterocytes that co-express *E-cadRNAi* and GFP were generated using MARCM (see Methods). **i,** If EGFR activation involves direct E-cadEGFR binding, then dpERK^+^ cells (red) will typically contact an *E-cadRNAi* enterocyte (green). If activation involves a dispersed signal, then dpERK^+^ cells will be close to, but not necessarily contact, an *E-cadRNAi* enterocyte. **j-k,** dpERK^+^ cells are enriched in the vicinity of a GFP^+^, *E-cadRNAi* enterocyte, but often do not contact. dpERK^+^ cells are rare in the vicinity of GFP^+^ control enterocytes. **l,** Spatial zone of EGFR activation. Each point is the measured distance between one dpERK^+^ cell and edge of the nearest *E-cadRNAi* enterocyte. dpERK^+^ cells frequentlylocalize 0-25 μm (~1-2 enterocyte diameters) away and can localize up to 50 µm away. N=4 midguts, analyzed 5 days after clone induction. One of three representative experiments is shown in each graph. Representative images are shown in all panels. All scale bars are 25 μm.

**Fig. 4.**
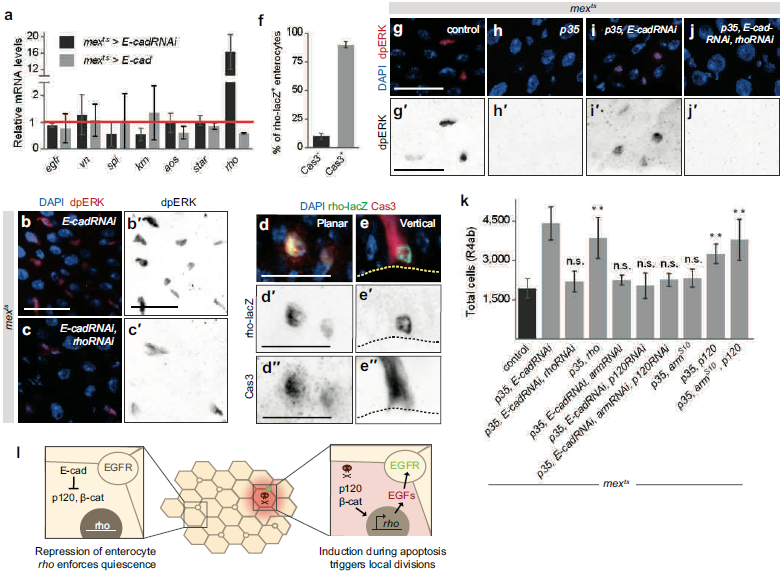
Enterocyte apoptosis activates stem cell division by disrupting E-cad-controlled inhibition of *rhomboid*. **a,** Enterocyte *E-cad* specifically inhibits expression of the obligate EGF protease *rhomboid* (*rho*). Levels of the indicated mRNAs were measured by qPCR of *mex*^*ts*^ control (red line), *mex*^*ts*^*>E-cadRNAi* (black bars), or *mex*^*ts*^*>E-cad* midguts (gray bars) after 4 days of induction. Relative to control, *rho* mRNAs increase by 15.5-fold upon *E-cad* depletion and decrease by 0.4-fold upon *E-cad* overexpression. mRNAs are not significantly altered for other components of EGF signaling: *egfr*; the EGF ligands *vein* (*vn*), *spitz* (*spi*), and *keren* (*krn*); or the post-translational EGF regulators *argos* (*aos*) and *star*. Values are means ± S.D. from 3 independent experiments. **b-c,** *E-cad*-depleted enterocytes require *rho* to hyperactivate stem cell EGFR. dpERK^+^ cells are abundant in *mex*^*ts*^*>E-cadRNAi* midguts but are substantially reduced in *mex*^*ts*^*>E-cadRNAi, rhoRNAi* midguts. Representative images are shown from two independent experiments; N=4 midguts per genotype in each experiment, analyzed after 2 days of transgene expression. See also Extended Data Fig. 3b. **d-f,** Expression of *rho* is activated in enterocytes during physiological apoptosis. Under steady-state conditions, the *rho-lacZ* reporter (green) is typically expressed in apoptotic enterocytes (red, cleaved caspase-3 staining) and rarely in non-apoptotic enterocytes. Planar (**d**) and vertical (**e**) views of two different fields are shown. In **e**, dotted line marks the basal epithelium. **f,** Quantification. Nearly all enterocytes that express *rholacZ* (90%) are also apoptotic. Values are means ± S.D from 3 independent experiments. N=3-4 midguts per experiment, analyzed 6 days post-eclosion; n=188 enterocytes total. **g-j,** Apoptosis-blocked enterocytes inhibit stem cell ERK activation via *E-cad* and *rho*. Compared to their nor-Mmal frequency, dpERK^+^ cells are strongly reduced when enterocyte apoptosis is blocked (*mex*^*ts*^*>p35)*, are restored when *E-cad* is additionally depleted (*mex*^*ts*^*>p35, E-cadRNAi*), and are strongly reduced again when both *E-cad* and *rho* are depleted (*mex*^*ts*^*>p35, E-cadRNAi, rhoRNAi*). Representative images are shown from two independent experiments; N=4 midguts per genotype in each experiment, analyzed after 2 days of transgene expression. See also Extended Data Fig. 3b. **k,** Activation of *rho* by p120-catenin and Armadillo drives organ hyperplasia. In apoptosis-inhibited midguts, loss of *E-cad* (*mex*^*ts*^*>p35, E-cadRNAi*) causes total cell number to increase by 128% compared to control, producing organ hyperplasia. Additional loss of *rho* (*mex*^*ts*^*>p35, E-cadRNAi, rhoRNAi*) restores normal cell number and prevents hyperplasia. On the other hand, overexpression of *rho* alone (*mex*^*ts*^*>p35, rho*) causes a 100% increase in total cells (*p*=0.0017), resulting in hyperplasia without loss of *E-cad*. Thus, *rho* is necessary and sufficient for hyper-plasia. Activation of *rho* is mediated by the E-cad-associated transcription factors p120-catenin (p120) and Armadillo (Arm) (Extended Data Fig. 7). Loss of either *p120* or *arm*, or both *p120* and *arm* (*mex*^*ts*^*>p35, E-cadRNAi, p120RNAi* and/or *armRNAi*), restores normal cell number and prevents hyperplasia. Overexpression of *p120*, but not constitutively active *arm* (*arm*^*S10*^), causes hyperplasia (69% increase in total cells; *p*=0.0011); overexpression of both *p120* and *arm*^*S10*^ (*mex*^*ts*^*>p35, p120, arm*^*S10*^) slightly exacerbates hyperplasia compared to *p120* alone (96% in-crease in total cells; *p*=0.0021). Thus, *p120* and *arm* are necessary and sufficient for hyperplasia. Values are means ± S.D from one of three representative experiments. *p* values (unpaired t-test) relative to control. N=4 midguts per genotype, analyzed after 4 days of transgene expression. See also Extended Data Fig. 3a. **l,** Model for homeostatic coupling of enterocyte apoptosis and stem cell division. In the absence of apoptosis (left), stem cells are quiescent because enterocyte E-cad represses p120- and Arm-dependent expression of *rho* to preclude activation of stem cell EGFR. Apoptotic enterocytes (right) disrupt this inhibitory feedback to trigger localized EGFR activation and replacement divisions of stem cells. Representative images shown in all panels. All scale bars are 25 μm.

How is enterocyte apoptosis coupled to stem cell divisions? The epithelial cell-cell adhesion protein E-cadherin (E-cad, also known as *shotgun*) drew our attention because it is a potential link between apoptosis, proliferation, and tissue homeostasis: First, E-cad undergoes targeted degradation by effector caspases in apoptotic epithelial cells ^23-26^. Second, loss of E-cad drives proliferation of epithelial tumors ^27^, and, in mouse intestine, targeted loss of enterocyte cadherin results in activated proliferation of progenitor cells ^28^. Third, the E-cad adhesion complex is an upstream regulator of density-dependent contact inhibition in cultured epithelial cells ^29-31^.

To assess whether E-cad is involved in coupling divisions to apoptosis, we first examined whether E-cad is downregulated in dying enterocytes. An E-cad::mTomato fusion ^32^ strongly delineated cell-cell interfaces between healthy enterocytes (Fig. 2a). In contrast, E-cad::mTomato was largely absent from cell-cell interfaces between dying enterocytes, which were identified by Sytox ^33^. Altogether, these findings imply that dying enterocytes lose cell-surface E-cad, akin to apoptotic cells in other epithelia ^23-26^.

To investigate the functional role of E-cad downregulation, we depleted *E-cad* in apoptosis-blocked enterocytes and measured stem cell divisions (Fig. 2b-e). In contrast to apoptotic inhibition alone (*mex*^ts^>*p35*), stem cells did not slow their divisions when *E-cad* was additionally knocked down (*mex*^ts^>*p35, E-cadRNAi*). At the same time, total cell number increased by 70%, resulting in enlarged, hyperplastic organs (Fig. 2h-j, Extended Data Fig 3a). These effects are *E-cad*-specific because depletion of another midgut cell-cell adhesion protein, *echinoid*, did not affect cell number (Fig. 2h). *E-cad* depletion alone (*mex*^ts^*>E-cadRNAi*) induced excess divisions but not organ hyperplasia, likely because of other, tissue-level effects (Fig. 2b, g; Extended Data Fig 5). *E-cad* overexpression suppressed divisions (Fig. 2b, f). Importantly, *E-cad* depletion did not disrupt the overall architecture or polarity of the midgut epithelium (Extended Data Fig. 4a, c-d, f), consistent with *E-cad* loss-of-function in the embryonic midgut ^34^; nor did it compromise the intestinal barrier, likely because septate junctions are intact (Extended Data Fig 4g-j). Thus, enterocyte E-cad suppresses stem cell divisions during apoptotic inhibition for homeostatic control of cell number.

E-cad mediates cell-cell adhesion by forming intercellular homodimers. We thus considered whether enterocyte E-cad acts by dimerizing with stem cell E-cad ^35,36^. To separately test the requirement for E-cad on progenitors, we built a system similar to Fig. 1f that combines genetic manipulation of stem and enteroblast cells (*esgGAL4;GAL80*^*ts*^, or *esg*^*ts*^) with split-*lacZ* lineage tracing of stem cells. Consistent with a prior report of 3-day *E-cad* null clones ^36^, we found that neither depletion nor overexpression of *E-cad* altered the rate of stem cell divisions in 4-day clones (Fig. 2k). These data suggest that enterocyte E-cad acts not in conjunction with stem cell E-cad, but rather via a distinct intermediary.

Prime candidates for this intermediary signal include Wingless/Wnt, Hippo, cytokine-JAK-STAT, and EGFR. These pathways act downstream of E-cadherin in other tissues ^30,37-42^ and are key mediators of injury-activated proliferation in the midgut ^43-45^. To assess whether these pathways are downstream of enterocyte E-cad, we examined whether *E-cad* depletion resulted in pathway activation. Known target mRNAs for Wingless and Hippo were not elevated in *E-cad* knockdown midguts compared to controls (Extended Data Fig. 6a). The cytokines *unpaired1-3* (*upd1-3*), which—particularly *upd3*—acutely respond to midgut injury ^46-49^, were not elevated (Extended Data Fig. 6a-d). Activation of downstream STAT targets in progenitors was also unaffected (Extended Data Fig. 6a, e-g). By contrast, EGFR target mRNAs were significantly elevated (Extended Data Fig. 6a, h-j).

To visualize EGFR activation, we immunostained midguts for the activated, diphosphorylated form of the EGFR effector ERK (dpERK) (Fig. 3a-f, Extended Data Fig. 3b). ERK activation occurred predominantly in stem cells (Fig. 3f), consistent with prior reports from others^44,46,50-54^. We found that ERK-activated stem cells were infrequent during normal turnover, became abundant when *E-cad* was depleted from enterocytes, and virtually disappeared when *E-cad* was overexpressed (Fig. 3a-c, Extended Data Fig. 3b). Other studies have reported that ERK activation in the midgut is predominantly due to activation of EGFR ^13,44,46,51-53^. Indeed, pharmacological EGFR inhibition (AG1478) or a conditional lethal heteroallele (*egfr*^*tsla*^*/egfr*^*f*^^24^) eliminated the dpERK signal in *E-cad* knockdown midguts (Fig. 3d-e, Extended Data Fig. 3b). Critically, EGFR signaling was required for excess stem cell divisions and organ hyperplasia caused by loss of *E-cad* in apoptosis-blocked enterocytes (Fig. 3g-h; Extended Data Fig. 3a). Thus, enterocyte E-cad inhibits stem cell EGFR signaling, and this inhibition mediates homeostatic control of cell number and organ size.

How does E-cad on enterocytes control EGFR activation in stem cells? Physical binding of E-cad and EGFR is one possible mechanism ^23-26,55,56^; dispersal of secreted signals is another ^57,58^. To shed light on these possibilities, we used MARCM ^59^ to generate single, GFP-marked enterocytes that were depleted of *E-cad.* We measured the spatial range of EGFR activation surrounding the marked enterocyte (Fig 3i). ERK-activated cells frequently appeared in the vicinity of, but did not necessarily contact, single *E-cad* knockdown enterocytes, supporting the existence of a dispersed signal. Strong EGFR activation occurred within a radius of ~25 μm from the edge of *E-cad* knockdown enterocytes, while weaker activation occurred up to ~50 μm (Fig. 3j-l). These findings suggest that EGFR activation involves a dispersed signal that acts in a localized zone around the enterocyte.

Is this dispersed, E-cad-controlled signal an EGF ligand? Supporting this notion, we found that the two enterocyte-derived EGFs, *spitz* (*spi*) and *keren* (*krn*) ^46,51,52^, were necessary for organ hyperplasia caused by loss of *E-cad* in apoptosis-blocked enterocytes (Figure 3h, Extended Data Fig. 3a). However, depletion of enterocyte *E-cad* surprisingly did not alter mRNA levels of either *spi* or *krn* (Fig. 4a). Furthermore, mRNA levels were unchanged for the visceral muscle-derived EGF *vein* ^46,51,52^, the EGF chaperone *star*, the secreted EGF inhibitor *argos*, and *egfr* itself. Strikingly by contrast, levels of the obligate EGF protease *rhomboid* (*rho*) increased substantially with *E-cad* knockdown and decreased with *E-cad* overexpression (Fig. 4a).

During EGF biosynthesis, Rho cleaves inactive, membrane-tethered EGF peptides into active, soluble forms ^60,61^. This function raises the possibility that E-cad controls EGF signaling by controlling EGF processing through Rho. Consistent with this possibility, we found that expression of *rho-lacZ* in enterocytes, but not diploid progenitor cells, was activated by *E-cad* knockdown and inhibited by *E-cad* overexpression (Extended Data Fig. 7c-d, f). Thus, E-cad suppresses transcription of *rho* specifically in enterocytes.

We tested whether levels of enterocyte *rho* determine levels of stem cell EGFR activation. Overexpression of *rho* in enterocytes promoted activation of ERK and increased numbers of mitotic stem cells (Extended Data Fig. 3b, Extended Data Fig. 7m-n,s). Conversely, depletion of *rho* abrogated activation of ERK (Extended Data Fig. 3b, Extended Data Fig. 7o). Furthermore, combined depletion of *rho* and *E-cad* together precluded the hyperactivation of ERK caused by depletion of *E-cad* alone (Fig. 4b-c, Extended Data Fig. 3b). Altogether, these results show that enterocyte E-cad inhibits stem cell EGFR by repressing enterocyte *rho*.

Is this E-cad-Rho-EGFR relay responsible for coupling stem cell divisions to enterocyte apoptosis? If so, then: (1) apoptotic enterocytes, which lose E-cad (Fig. 2a), should concomitantly upregulate *rho*; (2) loss of E-cad in apoptotic enterocytes should underlie stem cell EGFR activation; and (3) exogenous manipulation of *rho* should disrupt cellular equilibrium and alter organ size. We investigated each of these predictions. First, we examined the expression pattern of *rho* during normal midgut turnover. Strikingly, the *rho-lacZ* reporter predominantly marked apoptotic enterocytes and rarely marked non-apoptotic enterocytes (Fig. 4d-f). Thus, enterocytes repress *rho* when healthy but activate *rho* upon physiological apoptosis.

Prior studies have shown that *rho* is upregulated upon tissue-wide injury or pan-enterocyte death ^46,52^. Given this precedent, we wondered whether other injury signals are activated upon physiological apoptosis. However, the cardinal injury signal *upd3* was rarely observed in apoptotic enterocytes (Extended Data Fig. 6k). Furthermore, *upd3* was dispensible for stem cell ERK hyperactivation in *E-cad* knockdown enterocytes (Extended Data Fig. 3b, Extended Data Fig. 7i). These contrasts between *upd3* and *rho* indicate that dying cells signal differently in injury and steady-state contexts, possibly due to loss of the intestinal barrier or inefficient clearance of cell corpses following extensive damage.

To test the second prediction, we blocked enterocyte apoptosis (*mex*^*ts*^>*p35*) and examined stem cell EGFR activation. ERK-activated stem cells were virtually absent following apoptotic inhibition but were restored by the additional depletion of enterocyte *E-cad* (*mex*^*ts*^>*p35, E-cadRNAi*) in a *rho*-dependent manner (*mex*^*ts*^*>p35, E-cadRNAi, rhoRNAi*) (Fig. 4g-j, Extended Data Fig. 3b). These results demonstrate that loss of E-cad in apoptotic enterocytes is responsible for EGFR activation in stem cells.

To examine the third prediction, we manipulated *rho* in enterocytes and measured total cell number and organ size. Overexpression of *rho* in apoptosis-inhibited enterocytes (*mex*^*ts*^>*p35, rho*) resulted in organ hyperplasia, with cell number increased by 100% (Fig 4k, Extended Data Fig. 3a). Conversely, loss of *rho* in apoptosis-competent enterocytes (*mex^ts^>rhoRNAi*) resulted in organ atrophy, with cell number reduced by ~60% (Extended Data Fig. 8). This requirement for Rho during steady-state turnover contrasts with prior findings that Rho is dispensible for injury repair ^46^, drawing further distinction between turnover and repair mechanisms. Moreover, combined loss of both *rho* and *E-cad* in apoptosis-inhibited enterocytes (*mex*^*ts*^*>p35,E-cadRNAi, rhoRNAi*) thwarted the hyperplasia that would have resulted from loss of *E-cad* alone, and normal cell number was preserved (Fig. 4k, Extended Data Fig. 3a). Altogether, these results show that downstream of E-cad, *rho* is the pivot point that balances division and death to maintain constant organ size.

Finally, how does E-cad, a transmembrane adhesion receptor, control expression of *rho* in the nucleus? To address this question, we examined three factors whose ability to activate nuclear transcription is precluded by sequestration at E-cad-containing adherens junctions: β-catenin/Armadillo (Arm), p120-catenin (p120, also known as p120ctn), and YAP/Yorkie (Yki) ^38,62-68^. We found that *arm* and *p120*, but not *yki*, were required in *E-cad* knockdown enterocytes for both induction of *rho* and hyperactivation of stem cell EGFR (Extended Data Fig. 7a, g-l). Overexpression of *p120*, but not constitutively active *arm*^*S10*^, was sufficient for induction of *rho* (Extended Data Fig. 7b-f) and hyperactivation of EGFR (Extended Data Fig. 3b, Extended Data Fig. 7p-r). Overexpression of the transcriptional co-repressor Groucho, which can dimerize with β-catenin/Arm to repress *rho* in some tissues^69-71^, did not inhibit induction of *rho* in enterocytes (Extended Data Fig. 7a). Overexpression of the JNK inhibitor *puckered* partially inhibited *rho* induction, although the statistical significance of this effect was unclear (Extended Data Fig. 7a).

We next asked whether Arm, p120, or both affect organ size. Enterocyte knockdown of either *arm* or *p120* blocked the hyperplasia that otherwise would have occurred upon loss of *E-cad* and apoptotic inhibition; knockdown of both factors had a quantitatively similar effect (Fig 4k, Extended Data Fig. 3a). In addition, overexpression of *p120*, but not *arm*^*S10*^, was sufficient to induce hyperplasia, and overexpression of both factors exacerbated the effect (Fig. 4k, Extended Data Fig 3a). Conversely, depletion of either *arm* or *p120* produced mild atrophy (Extended Data Figure 8). These data show that the p120 and Arm transcription factors underlie Ecad-controlled expression of *rho* and suggest that E-cad represses *rho* by sequestering p120 and Arm to control organ size.

Altogether, our results demonstrate that steady-state organ turnover is not driven by the constitutive cycling of stem cells. Instead, healthy enterocytes keep stem cells in a default state of quiescence, while the sporadic appearance of apoptotic enterocytes triggers replacement divisions. Precise cellular balance hinges upon the E-cad-dependent repression of *rho* in healthy enterocytes, which is disrupted when E-cad is lost in apoptotic enterocytes (Fig 4l). Because divisions are coupled to apoptosis, turnover remains zero-sum over time.

The direct coupling of divisions to apoptosis suggests a simple explanation for how the midgut epithelium dynamically maintains a constant number of cells with such robust precision. Crucially, a single midgut enterocyte can efficiently activate EGFR in stem cells within a ~25 μm radius (Fig. 3j-l). We suggest that this zone of activation, which exists only for as long as the dying cell remains in the epithelium, may be critical for homeostatic size control. If, by chance, stem cells produce too many enterocytes, then the stem cells’ physical spacing would increase; subsequently, fewer stem cells would be within the activation zone of the next dying enterocyte, and fewer divisions would result. Similarly, too few enterocytes would place more stem cells in the activation zone, and more divisions would result. By setting the steady-state number of enterocytes, the integration of these activation zones over the entire epithelium would determine overall organ size. In this manner, localized cell-cell communication can give rise to tissue-level homeostatic equilibrium.

Our study brings to light a basic distinction in how EGFs are deployed during steadystate turnover versus injury repair. At steady-state, apoptotic induction of *rho* is strictly cell autonomous because E-cad-dependent activation of p120 and Arm is confined to the dying enterocyte. This cell autonomous pathway limits the release of EGFs to the precise time and place that a new cell is needed, as appropriate for zero-sum replacement. By contrast during injury, induction of *rho* and EGFs involves an additional, non-cell autonomous pathway in which damaged enterocytes upregulate *upd3*; Upd3 in turn activates enteroblasts and visceral muscle to upregulate *rho* and *EGF*s ^46,48,49,52^. This non-autonomous pathway permits EGFs to be released in a widespread, indiscriminate manner, as appropriate for an emergency response. Underscoring this distinction, enterocyte *upd3* is required for repair ^13,46,48,49^ but not homeostasis (Extended Data Figs. 6a-d, k and 7i), whereas enterocyte *rho* is required for homeostasis (Fig 4) but not repair ^46^.

Stem cell EGFR signaling is known to affect homeostasis of other tissues ^72-74^, raising the possibility that spatially specific control of EGFR activation by E-cad and Rho is a general mechanism for cellular equilibrium. By extension, loss of spatial control should lead to pathological loss of homeostasis. Indeed, we note that multiple human carcinomas downregulate E-cad, upregulate Rhomboids, and activate EGFR ^27,75-77^, and that progression of colorectal carcinoma, which initiates through loss of the catenin-destabilizing factor *APC* (adenomatous polyposis coli), requires upregulation of the mammalian Rhomboid *RHBDD1* ^78^. Given these intriguing links, we propose that insights into the development of epithelial cancers may emerge from understanding E-cad-EGFR feedback control of steady-state epithelial turnover.

## METHODS

### *Drosophila* Husbandry

Crosses utilizing the GAL4/GAL80^ts^ system were performed at 18°C. Upon eclosion, adult animals remained at 18°C for 4 days, unless otherwise indicated. On adult day 4, animals were temperature shifted to 29°C to inactivate GAL80^ts^ and induce GAL4-mediated expression. Midguts were harvested for immunostaining after appropriate lengths of induction (see figure legends for individual experiments). All other crosses were performed at 25°C; refer to figure legends for individual timepoint information. Adult female flies were used in all experiments.

### Fly Stocks

*w; esgGAL4, tubGAL80ts, UAS-GFP; UAS-flp, act<CD2<GAL4* (*esg*^*F/O*^) (Bruce Edgar) ^13^

*mexGAL4* (Carl Thummel)

*esgGAL4* (Ben Ohlstein)

*y, w; TI{TI}shg[mTomato]* (Bloomington)

UAS-*E-cadherinRNAi* (TRiP.HMS00693) (Bloomington)

UAS-*E-cadherinRNAi* (TRiP. JF02769) (Bloomington)

UAS-*E-cadherinRNAi* (TRiP.GL00646) (Bloomington)

UAS-*echinoidRNAi* (TRiP.GL00648) (Bloomington)

UAS-*rhomboidRNAi* (TRiP.JF03106) (Bloomington)

UAS-*spitzRNAi* (TRiP.HMS01120) (Bloomington)

UAS-*kerenRNAi* (KK104299) (VDRC)

UAS-*armadilloRNAi* (TRiP.JF01251) (Bloomington)

UAS-*armadilloRNAi* (KK107344) (VDRC)

UAS-*p120ctnRNAi* (TRiP.HMC03276) (Bloomington)

UAS-*yorkieRNAi* (TRiP.JF03119) (Bloomington)

UAS-*unpaired3RNAi* (TRiP.HM05061) (Bloomington)

UAS-*hisH2A:RFP* (Bloomington)

UAS-*p35* (Bloomington) UAS-*diap1* (Bloomington)

UAS-*E-cadherin*^*DEFL*^ (Margaret Fuller) ^79^

UAS-*rhomboid* (Bloomington) UAS-*armadillo*^*S10*^ (Bloomington)

UAS-*p120ctn* (Bloomington) UAS-*groucho* (Amir Orian)

UAS-*puckered* (*puc2A*) (Huaqi Jiang)

*y w hsflp; X-15-29 w*^*+*^ (‘split-*lacZ*’) ^19^

*y w; y*^*+*^ *X-15-33* (‘split-*lacZ*’) ^19^

*Egfr*^*f24*^*/T(2;3)TSTL* (Bloomington) *Egfr*^*tsla*^*/T(2;3)TSTL* (Bloomington)

*w UAS-CD8:GFP hsflp; tubGAL4; FRT82 tubGAL80* (David Bilder) ^6^

*w; FRT82* (David Bilder) ^6^

*rho*^*X81*^ (*rho-lacZ*) (Huaqi Jiang) ^80^

*10xSTAT-GFP* (Bloomington)

*Upd3.1-lacZ* (Huaqi Jiang)

*cycE-lacZ* (Bloomington)

Detailed information on *Drosophila* genes and stocks is available from FlyBase (http://flybase.org/).

### Immunohistochemistry and Microscopy

Samples were fixed, immunostained, and mounted as previously described ^6^. Primary antibodies: mouse anti-β-galactosidase (1:400, Promega Z3781), mouse anti-Armadillo (1:100, DSHB N27A1), rabbit anti-cleaved caspase 3 (1:200, Cell Signaling, generous gift from D. Bilder ^6^) rabbit anti-diphospho-ERK (1:400, Cell Signaling 4370P), goat anti-HRP-Cy3 (Cappel, 1:100) which stains stem cells and enteroblasts ^6^, mouse anti-Coracle (1:50, DSHB C615.16), mouse anti-Discs large C615.16 (1:50, DSHB 4F3), and rabbit anti-phospho-histone H3, Ser 10 (1:1000, EMD Millipore). Secondary antibodies: Alexa Fluor 488-, 555- or 647-conjugated donkey anti-rabbit or anti-mouse IgGs (1:800, LifeTechnologies A31570, A11001, and A21244). Nuclei were stained with DAPI (LifeTechnologies, 1:1000). Actin was stained with SiR-Actin (Spirochrome, 1:500) or Alexa 647-conjugated phalloidin (1:100, LifeTechnologies). Samples were mounted in ProLong (LifeTechnologies). Imaging of samples was performed on a Leica SP8 confocal microscope, with serial optical sections taken at 3.5 μm intervals through the entirety of whole-mounted, immunostained midguts.

### Regionalization of the Adult Midgut; Cell Counts and Size Measurements of the R4ab (P1-2) compartment

The *Drosophila* midgut is compartmentalized along its proximal-distal axis. Each compartment exhibits a characteristic digestive physiology, gene expression pattern, and stem cell division rate ^14,15,81,82^. In general, stem cell clones do not cross compartment boundaries ^15^. Our study focused specifically on two adjacent compartments, known alternatively as R4ab or P1-2, which comprise the major region of nutrient absorption ^14,15^. We observed that R4ab consistently exhibited complete cellular turnover between adult days 4-8, as indicated by *esg*^*F/O*^ labeling (Fig. 1a-e, Extended Data Fig. 1f-g). Other midgut compartments exhibited variable, incomplete turnover during the same time period, consistent with prior reports ^9,83-85^; they were not analyzed in this study.

To perform total cell counts of R4ab, this region was first identified in confocal image stacks using morphological landmarks (Extended Data Fig 1b-e, g)^14^ and digitally isolated in Fiji. Bitplane Imaris software algorithms were applied to generate three-dimensional organ reconstructions and comprehensively count individual cell nuclei by mapping DAPI signals to Imaris surface objects. For analysis of *esg*^*F/O*^ midguts, GFP^+^ cells were additionally counted by mapping DAPI/GFP colocalization signals to Imaris surface objects. R4ab lengths were measured by a spline through the center of individual midguts in Fiji.

### Split*-lacZ* Clone Induction and Analysis

As depicted in Fig. 1f, animals were raised at 18°C and shifted to 29°C four days post-eclosion. Split*-lacZ* clone induction ^19^ was performed by subjecting animals to two 30-min, 38.5°C heat shocks separated by a 5-min chill on ice. Four days after clone induction, midguts were immunostained and clones in the R4ab region were identified and analyzed by visual examination of serial confocal sections. Clones in regions outside R4ab were excluded from analysis. Clone size was measured as the number of contiguous cells in one discrete clone, as previously described ^6^. Approximately 1-3 single labeled enterocytes per R4ab region were observed, consistent with published reports ^9,86^; these transient clones were not included in our quantification. Single labeled diploid cells were included because these likely represented individual stem cells that did not divide during the chase period. No labeled cells were observed in the absence of 38.5°C heat shock.

#### Safeguards to ensure exclusion of non-stem cell (transient) clones

To ensure that clone counts comprised exclusively stem cell clones and excluded any non-stem cell (transient) clones that were directly labeled by the heat shock, our split*-lacZ* clonal analyses incorporated two, redundant safeguards. First, a 4-day chase period was included between heat-shock induction and subsequent clonal analysis. Because they are post-mitotic and transient, enteroblasts/enterocytes that were directly labeled by the heat shock would have been lost during the succeeding chase period. Confirming that transient clones were nearly absent, only 1-3 single labeled enterocytes were observed per midgut R4ab region after the 4-day chase. As a second safeguard, all single, labeled enterocytes were excluded from our clone counts.

### Sytox staining

Sytox Green (ThermoFisher, 5mM in DMSO) or Sytox Orange (ThermoFisher, 5mM in DMSO) were diluted 1:5,000 in 5% sucrose. Sytox solution was fed to animals in an empty vial for 5-6 hours, after which midguts were dissected and mounted in ProLong (LifeTechnologies). Because Sytox is incompatible with fixation, live organs were imaged immediately after mounting.

### MARCM Clone Inductions

MARCM clone inductions ^59^ were performed by subjecting animals to two 30-min, 38.5°C heat shocks separated by a 5-min chill on ice. For single-enterocyte MARCM clones, animals were dissected five days post-induction and terminal clones consisting of one GFP^+^ enterocyte (identified by its polyploid nucleus) were selected for analysis. GFP^+^ enterocytes were excluded from analysis if another GFP^+^ clone was present within an 80 μm radius. Fiji was used to measure the distance between the plasma membrane of the nearest GFP^+^ enterocyte and the center of dpERK^+^ stem cells within a 60 μm radius. For mosaic analyses of multicell MARCM clones, animals were fed Sytox three days post-induction and dissected. The proportion of labeled clone cells (GFP^+^) that were also Sytox^+^ was quantified.

### AG1478 Drug Treatment

Stocks of AG1478 (Sigma) were dissolved in EtOH and subsequently diluted in dH_2_O to reach a working concentration of 100 μM AG1478 (in 0.02% EtOH). This 100 μM stock solution was used to prepare yeast paste, which was fed to animals as a supplement to their standard cornmeal-molasses diet for the duration of induced gene expression.

### Smurf Assay

Smurf assays ^87^ were conducted by feeding adult animals yeast paste containing 2.5% Brilliant Blue FCF (Sigma) and scoring animals for leakage of dye into the abdomen. Animals were scored as ‘non-Smurf’ if the blue dye was confined to the GI tract and ‘Smurf’ if blue dye leaked outside the GI tract. As a positive control, animals were fed dye in conjunction with 1% SDS.

### qRT-PCR

mRNA was extracted from midguts (5 animals/experiment) followed by cDNA synthesis with Invitrogen SuperStrand III First Script Super Mix (Invitrogen). Real-time PCR was performed using the relative standard curve method with SYBR GreenER Supermix (Invitrogen) on a StepOnePlus ABI machine. Expression levels were normalized to *mexGAL4*^*ts*^*>CD4-GFP* midguts; *mef2* transcripts were used as a reference ^6^.

### Statistical Analysis

All statistical analysis was performed using Graphpad Prism 6. For comparisons of clone size distributions, unpaired two-tailed Mann-Whitney tests were used to assess statistical significance. (Clone size distributions are non-normal, independent, and derived from a simple random sample.) For comparisons of cell numbers and gut length, unpaired two-tailed t-tests were used to assess statistical significance. (Organ cell number and size distributions are normal, independent, and derived from a simple random sample.) For comparisons of *rho* gene expression, unpaired two-tailed t-tests were used to assess statistical significance. Legend: ns = not significant (*p*>0.05), *= *p*<0.05, **= *p*<0.01, ***= *p*<0.001, and **** = *p*<0.0001.

### Study Design

Sample sizes were chosen based on our previous study ^6^, which also characterized changes in organ cell number and clone sizes. In *split-lacZ* experiments, single enterocyte clones were excluded from analysis. No other exclusion criteria were applied. No sample randomization or blinding was performed, although automated, Imaris-based computer algorithms were used to analyze and quantify most data in this study.

### qPCR Primers

Primers for qPCR listed from 5’ to 3’:

*vein*-fwd GAACGCAGAGGTCACGAAGA

*vein*-rev GAGCGCACTATTAGCTCGGA

*spitz*-fwd CGCCCAAGAATGAAAGAGAG

*spitz*-rev AGGTATGCTGCTGGTGGAAC

*keren*-fwd CGTGTTTGGCAACAACAAGT

*keren*-rev TGTGGCAATGCAGTTTAAGG

*egfr*-fwd TGCATCGGCACTAAATCTCGG

*egfr*-rev GGAAGCTGAGGTCCAAATTCTC

*argos*-fwd TGCTGTTGGGTGAATTTCAGG

*argos*-rev CGACTGGTCCAGATGATCCA

*star*-fwd AGCCCAGTCCTTCAAACCC

*star*-rev CCACAGTCTTTGGTTGGTTGC

*rhomboid*-fwd GAGCACATCTACATGCAACGC

*rhomboid*-rev GGAGATCACTAGGATGAACCAGG

*frizzled* 3-fwd TCTTGTGCCCGCAAAACTTTA

*frizzled* 3-rev CCTAGAATGAGGGTCTCAGACG

*senseless*-fwd GATCGTGACTTTGCCTTGACG

*senseless*-rev CCTGATAGTCCTGCTTGCTGT

*expanded*-fwd GATGCTGGACACCGAACTCT

*expanded*-rev CTTGCTCTCGGGATCTGC

*diap1*-fwd GAAAAAGAGAAAAGCCGTCAAGT

*diap1*-rev TGTTTGCCTGACTCTTAATTTCTTC

*pointed*-fwd CTACGAGAAGCTGAGTCGCG

*pointed*-rev TATCGTTTGCCTGCCGTCTT

*cycE*-fwd ACAAATTTGGCCTGGGACTA

*cycE*-rev GGCCATAAGCACTTCGTCA

*upd1*-fwd CCTACTCGTCCTGCTCCTTG

*upd1*-rev TGCGATAGTCGATCCAGTTG

*upd2*-fwd GAGGGCAGCTACGACAGTG

*upd2*-rev GGAGAAGAGTCGCAGGTTGT

*upd3*-fwd AAATTCGACAAAGTCGCCTG

*upd3*-rev TTCCACTGGATTCCTGGTTC

*wdp*-fwd TGGCAACCACAATGAGGAACAG

*wdp*-rev GACCGAGAAGACCTTCCAGTCAAC

*Socs36E*-fwd CAGTCAGCAATATGTTGTCG

*Socs36E*-rev ACTTGCAGCATCGTCGCTTC

*mef2*-fwd ATCGGCAGGTGACCTTCAAC

*mef2*-rev GTTGTACTCGGTGTACTTGAGCAG

Primer sequences from Jiang et al. 2011 ^46^, Shaw et al. 2010 ^88^, and Fly Primer Bank (http://www.flyrnai.org/FlyPrimerBank).

### Genotypes by Figure

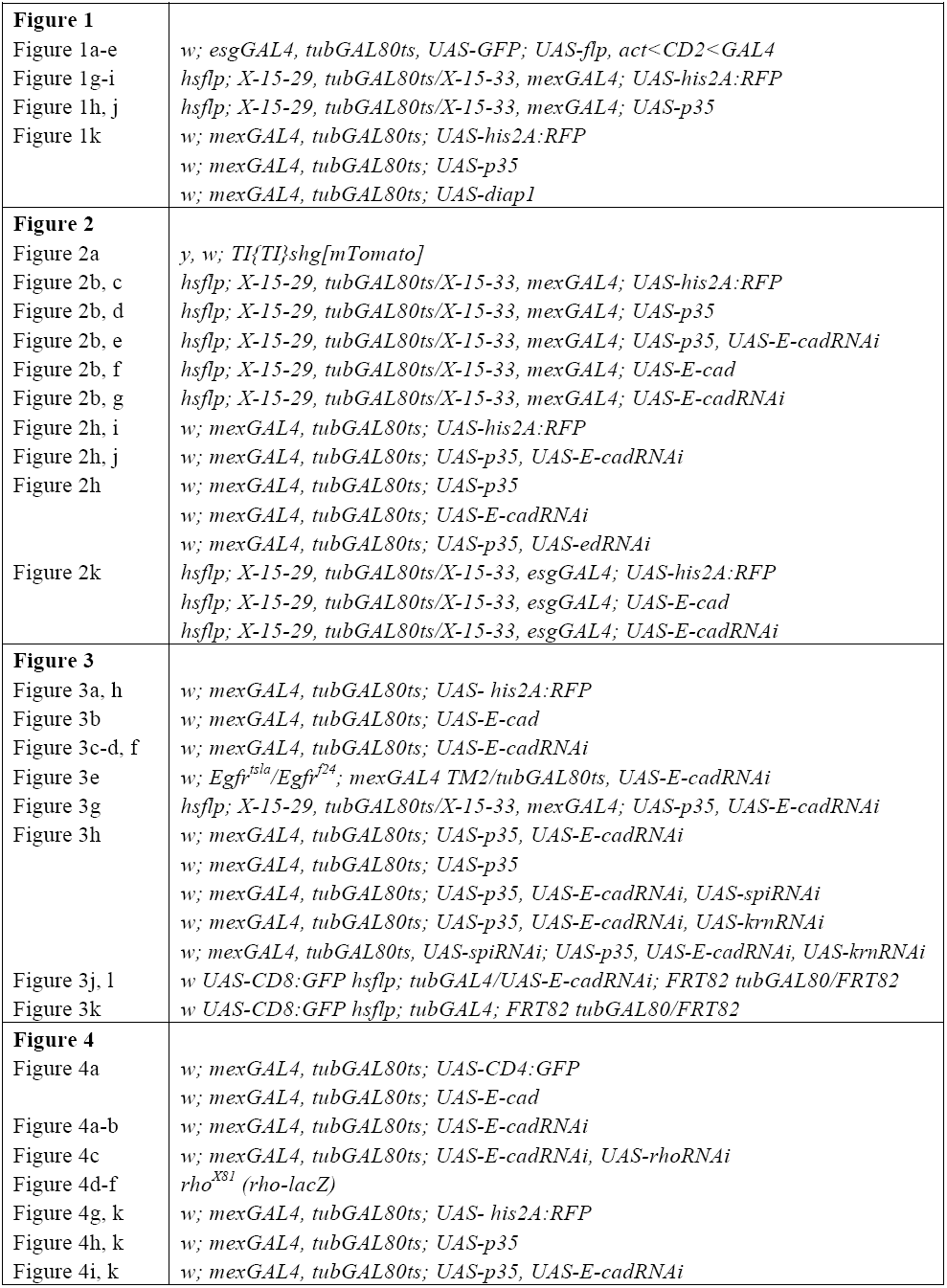
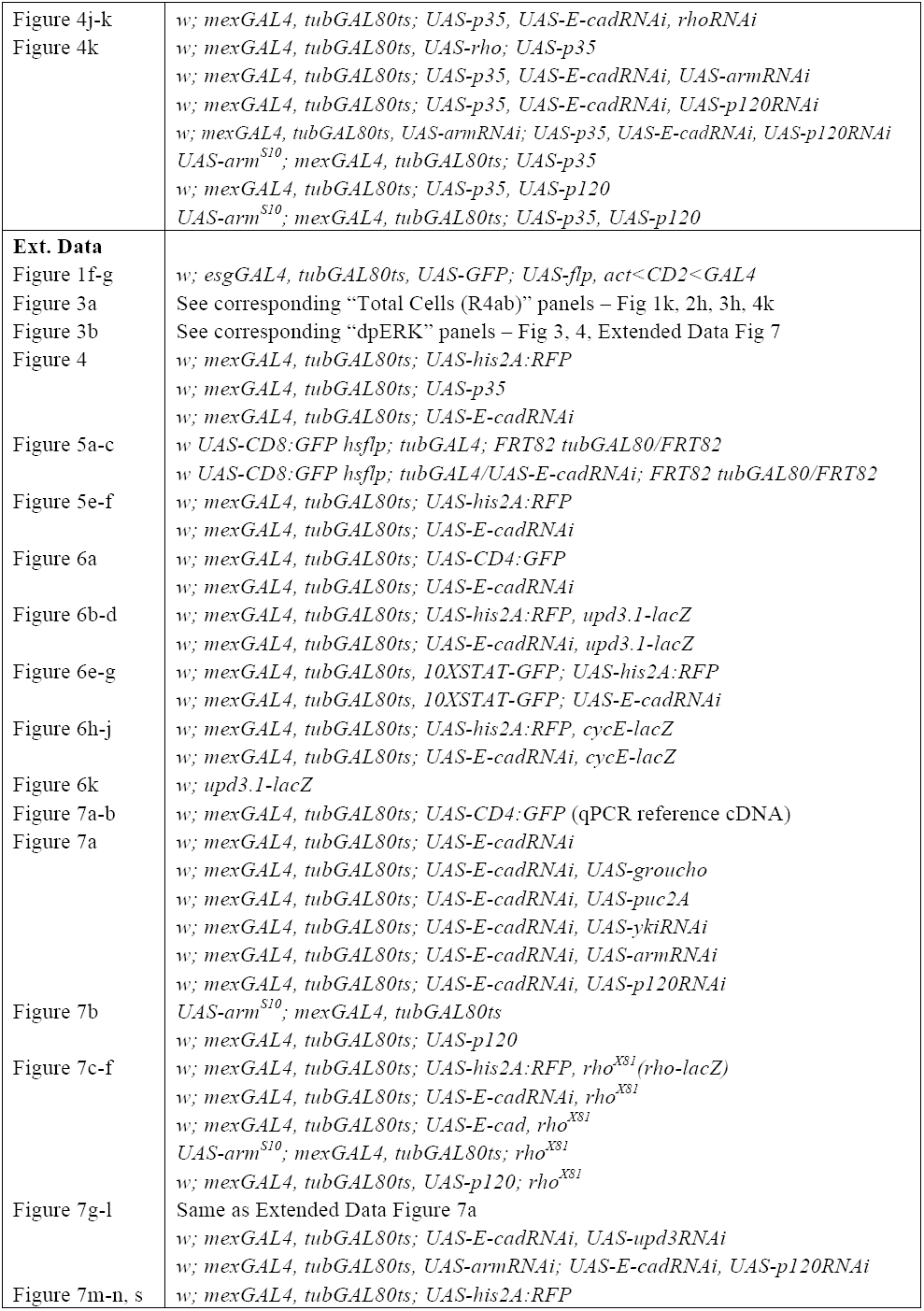
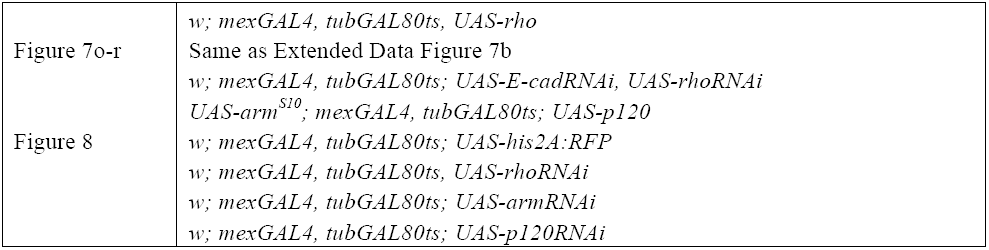

## Acknowledgments

J.L. was supported by NSF GRFP DGE-114747 and NIH T32GM007276. This work was supported by NIH R03DK104027 and R01GM116000-01A1 to L.E.O. Confocal microscopy was performed at the Stanford Beckman Cell Sciences Imaging Facility (NIH 1S10OD01058001A1). We thank D. Bilder for the gift of cCas-3 antibody; the Developmental Studies Hybridoma Bank for other antibodies; D. Bilder, B. Edgar, M. Fuller, H. Jiang, B. Ohlstein, C. Thummel, the Bloomington Drosophila Stock Center (NIH P40OD018537), the TRiP at Harvard Medical School (NIH/NIGMS R01-GM084947), and the Vienna Drosophila Resource Center for fly stocks; J. Axelrod, M. Goodman, M. Fuller, W.J. Nelson, R. Nusse, M. Krasnow,T. Nystul, and D. Fox for comments on the manuscript; and B. Benham-Pyle, M. Mirvis, N. Pierce, and D. Gordon for helpful discussions. The authors declare no competing interests.

## Author contributions

J.L. and L.E.O. designed the experiments and wrote the manuscript. J.L., S.B., and S.N. performed confocal microscopy. J.L. and S.N. prepared microscopy specimens. J.L. performed all other experiments, genetic crosses, data analysis, and statistical analysis.

## Author information

The authors declare no competing financial interests. Correspondence and requests for materials should be addressed to L.E.O.

## EXTENDED DATA FIGURES 1-8

**Extended Data Fig. 1.**
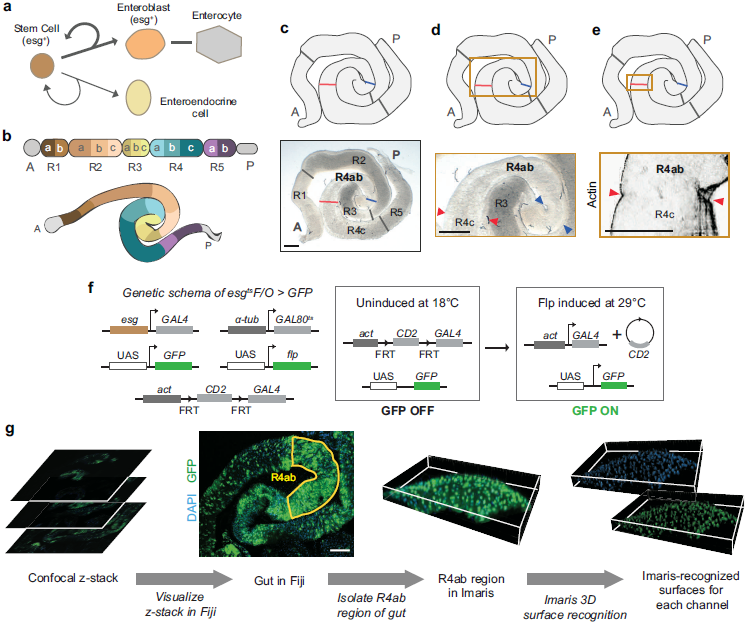
Midgut lineage and morphology, *esg* ^*F/O*^ labeling system, and workflow for semi-automated cell counting. **a,**Lineage of the adult *Drosophila* midgut ^9,84,89^. In general, stem cells are the only cells capable of division. Asymmetric stem cell divisions typically produce absorptive enterocytes; less frequently, they produce secretory enteroendocrine cells. Enterocytes arise through direct maturation of transient, post-mitotic intermediates called enteroblasts. Both stem and enteroblast cells express the Snail-family transcription factor *escargot* (*esg*). **b**, Compartments of the female adult midgut ^14,15,81^. R4ab was used for all experiments in this study. Schematic adapted from ^14,81^. **c-e**, Identification of R4ab through morphological landmarks. As defined in ^14^, R4ab is bounded by the apex of the midgut tube’s most distal 180° turn (blue arrowheads) and by the first prominent muscle constriction distal to this 180° turn (red arrowheads). **e**, The R4ab distal muscle constriction is particularly apparent in confocal optical sections ^14^. Visceral muscle stained with phalloidin. Midguts in panels **c**-**d** and **e** are two different samples. **f,** Genetic schema of the *esg*^F/O^ system ^13^. Stem and enteroblast cells are induced to express heritable GFP by temperature shift ts from 18°C to 29°C. The temperature shift inactivates GAL80, which allows *esgGAL4* to drive expression of both *UAS-GFP* and *UAS-flp* in stem and enteroblast cells. In these cells, flp recombinase renders GFP expression permanent and heritable by excising a CD2 ’flp-out’ cassette to generate a functional *actGAL4*; once generated, *actGAL4* drives expression of *UAS-GFP* (and *UAS-flp*) irrespective of cell type. Thus, after temperature shift, all mature cells that arise from undifferentiated cells will express *GFP*. **g,** Pipeline for semi-automated, comprehensive cell counts of 3D, reconstructed midgut regions. (1) Confocal microscope z-stacks capturing the entire depth of the organ are visualized in Fiji. (2) The R4ab region of the midgut (yellow outline) ^14,15^ is digitally isolated and exported to Imaris. (For illustrative purposes, only the top half of the gut tube is shown.) Note that different midgut regions have different rates of turnover: R4ab undergoes complete turnover between adult days 4-8 (at 29°C). However, other regions undergo slower turnover, as shown by large unlabeled regions outside of R4ab. The slower turnover of these other regions is consistent with the 7-21 day time frame of whole-organ turnover reported by others ^9,83-85^. See Methods for further discussion. (3) To quantify total cells (DAPI^+^), nuclei are mapped to surface objects using Imaris (Figs. 1e, 1k, 2h, 3h, 4k; Extended Data Fig. 8a). To quantify newly-added cells in the *esg*^F/O^ system (Fig. 1e), GFP^+^ nuclei are recognized in Imaris by co-localization of GFP and DAPI channels, and subsequently mapped to surface objects. Scale bars are 100μm.

**Extended Data Fig. 2.**
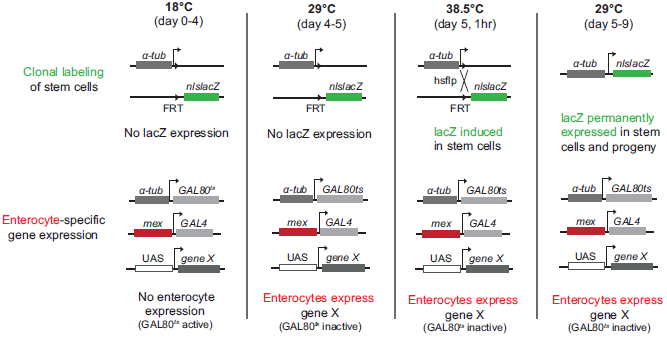
Genetic schema of system to simultaneously manipulate enterocyte expression and trace stem cell divisions. Detailed explanation of the genetic system in Fig. 1f. Animals are raised at 18°C; at this temperature, GAL80^ts^ represses *mex*-driven GAL4 in enterocytes, and lacZ labeling of stem cells is not induced. When animals are temperature-shifted to 29°C, consequent inactivation of GAL80^ts^ allows *mex*-driven GAL4 to express genes of interest (UAS-gene X) specifically in enterocytes. After 1 day of UAS gene expression, animals are shifted to 38.5°C for one hour to induce ubiquitous expression of *flp* recombinase, which is under control of a heat-shock promoter (*hs-flp*). Flp catalyzes trans-recombination of the two FRTs to place the *α-tubulin* promoter upstream of the promoter-less *nls:lacZ* cassette and, consequently, turn on permanent *nls:lacZ* expression. After heat shock, animals are returned to 29°C to maintain UAS-trangene expression. Midguts are harvested for clonal analysis 4 days after the 38.5°C heat shock. This 4-day chase, combined with exclusion of single, labeled enterocytes from clone counts, ensures that counts comprise exclusively stem cell clones and that any non-stem (transient) clones are eliminated (see Methods).

**Extended Data Fig. 3.**
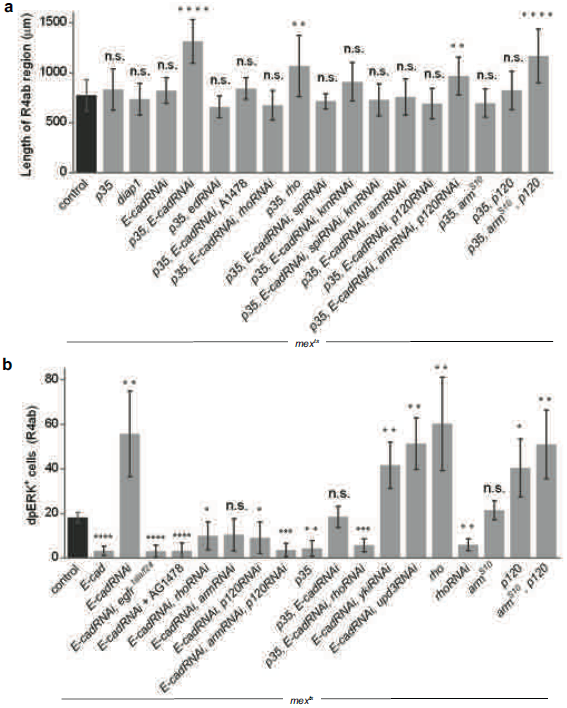
Quantifications of organ size and EGFR activation in genetically manipulated midguts. **a,** Lengths of the R4ab compartment. For 4 of 17 conditions, R4ab are significantly longer than control: *mex*^*ts*^ *>p35, E-cadRNAi* (unpaired t-test: *p*<0.0001), *p35, rho* (*p*=0.0052), *p35, E-cadRNAi, armRNAi, p120RNAi* (*p*=0.0029), and *p35, p120, arm*^*S10*^ (*p*<0.0001). N=10-12 midguts per genotype, analyzed after 4 days of UAS-transgene expression. Values are means ± S.D. **b**, Quantifications of dpERK^+^ cells in the R4ab compartment. For 15 of 18 conditions, numbers of dpERK^+ *ts*^ cells are significantly different from control: *mex >E-cad* (unpaired t-test: *p*<0.0001), *E-cadRNAi* (*p*=0.0081), *E-cadRNAi, egfr*^*tsla/f24*^ (*p*<0.0001), *E-cadRNAi* + AG1478 (*p*<0.0001), *E-cadRNAi, rhoRNAi* (*p*=0.04), *E-cadRNAi, p120RNAi* (*p*=0.04), *E-cadRNAi, armRNAi, p120RNAi* (*p*=0.007), *p35* (*p*=0.0013), *p35, E-cadRNAi, rhoRNAi* (*p*=0.0005), *E-cadRNAi, ykiRNAi* (*p*=0.006), *E-cadRNAi, upd3RNAi* (*p*=0.002), *rho* (*p*=0.007), *rhoRNAi* (*p*=0.001), *p120* (*p*=0.015), *arm*^*S10*^, *p120* (*p*=0.005). N=4 midguts per genotype, analyzed after 2 days of UAStransgene expression. Values are means ± S.D.

**Extended Data Fig. 4.**
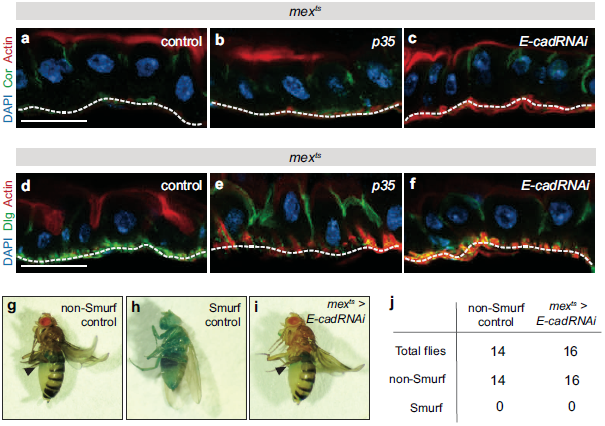
Analysis of epithelial architecture, polarity, and barrier function. **a-f,** Apoptotic inhibition or *E-cad* depletion in enterocytes does not disrupt epithelial architecture or apical-basal polarity. Images show vertical sections through the midgut epithelium after 4 days of either *mex* ^*ts*^ *>p35* or *mex* ^*ts*^ *>E-cadRNAi* expression. Enterocytes remain as a coherent monolayer. Apical-basal polarity is intact, as revealed by immunolocalization of apical, actinrich microvilli (**a-f**, red) and of apico-lateral septate junction proteins Coracle (**a-c**, green) and Discs-large (**d-f**, green). At the basal surface of the epithelium (white dotted lines), midgut visceral muscle cells stain brightly for actin and Discs-large. Actin stained with SiR-Actin. Scale bars are 25µm. **g-j,** Depletion of *E-cad* in enterocytes does not compromise the intestinal barrier. To test the intestinal barrier, animals were subjected to Smurf assays in which a blue, nonabsorbable food dye is administered by feeding ^87^. The dye remains within the midgut when the barrier is intact (**g**, non-Smurf) but leaks into the body cavity when the barrier is compromised, such as after consumption of 1% SDS (**h**, Smurf). After 10 days of *mex* ^*ts*^ *>E-cadRNAi* expression, midguts still retain the blue dye; no Smurf phenotypes are observed (**i-j**).

**Extended Data Fig. 5.**
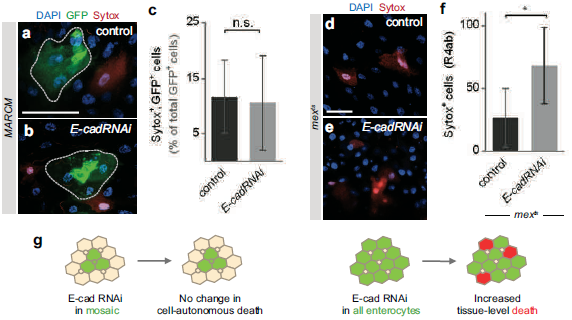
Depletion of *E-cad* has distinct cell-autonomous and tissue-level effects on cell death. The data in Fig. 2h show that midguts accumulate excess cells when *E-cad* is depleted from apoptosis-inhibited enterocytes but not apoptosis-competent enterocytes. To shed light on this difference, we examined whether *E-cad* depletion itself promotes cell death. Two approaches, mosaic knockdown and pan-enterocyte knockdown, were used to distinguish direct, cellautonomous effects from indirect, tissue-level effects. **a-c,** Mosaic knockdown of *E-cad* does not promote cell-autonomous death. Mosaic midguts are generated by using MARCM ^59^ to induce sparse, multicellular, GFP-marked clones in a background of unmarked, genetically unperturbed cells. **a-b,** Dotted outlines show representative control and *E-cadRNAi* clones (green). Sytox (red) identifies dying cells. **c**, Percentage of GFP^+^ cells that are also Sytox^+^. Dying cells occur with near-equal frequency within control and *E-cadRNAi* clones. Unpaired t-test, *p*>0.05. N=5 midguts per genotype, analyzed 9 days after clone induction; n=873 cells in control clones and 698 cells in *E-cadRNAi* clones. Values are means ± S.D. **d-f,** Pan-enterocyte knockdown of *E-cad* promotes cell death, likely through a non-autonomous effect. **d-e,** Representative images of *mex*^*ts*^ control and *mex*^*ts*^*>E-cadRNAi* epithelia. Sytox (red) identifies dying cells. **f,** Quantification of Sytox^+^ cells in the R4ab compartment. The number of dying cells increases ~2.5x in *E-cadRNAi* midguts compared to control (unpaired t-test, *p*=0.03). N=5 midguts per genotype, analyzed after 3 days of transgene induction. Values are means ± S.D. Scale bars are 25 μm. **g,** Summary. The unaltered frequency of dying cells in *E-cadRNAi* mosaic clones indicates that loss of *E-cad* does not cause cell-autonomous death. This result suggests that elevated death in *mex*^*ts*^*>E-cadRNAi* guts is a non-autonomous, tissue-level effect, possibly due to excess divisions (Fig 2b) and consequent crowding ^90^. These findings may explain why *p35, E-cadRNAi* guts accumulate excess cells whereas *E-cadRNAi* guts retain a normal number of cells (Fig. 2h).

**Extended Data Fig. 6.**
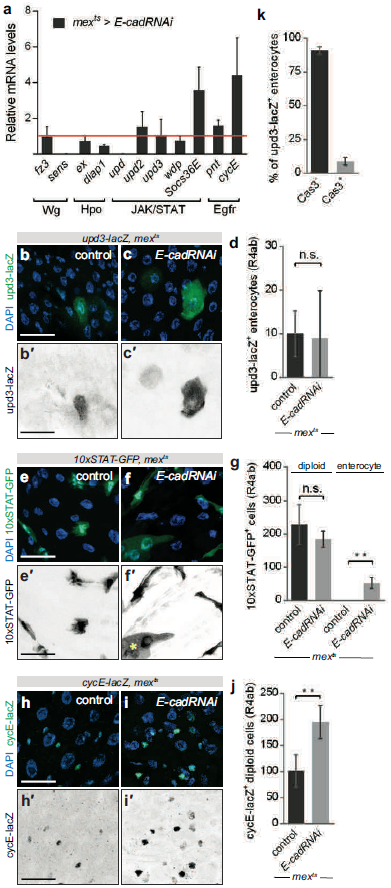
Loss of enterocyte *E-cad* activates EGFR, but not Wg, Hpo, or Upd/JAK/STAT. **a,** Effect of enterocyte *E-cad* depletion on target mRNAs of known midgut regulatory pathways. mRNAs were measured by qPCR of *mex* ^*ts*^ control or *mex*^*ts*^ *>E-cadRNAi* midguts. Relative to control (red line), mRNAs are unchanged for: the Wg targets *frizzled-3* (*fz3*) and *senseless* (*sens*) ^91,92^, the Hpo/Yki targets *expanded* (*ex*) and *diap1* ^88,93^, the injury-associated cytokines *upd* and *upd3* ^46-49^, and the JAK/STAT target *windpipe* (*wdp*) ^94^. The other JAK/STAT target, *Socs36E*, is elevated, likely reflecting its occasional activation in enterocytes (panel **f**). By comparison, the EGFR target *pointed* (*pnt*) ^52^ is slightly increased, and the EGFR target *cyclinE* (*cycE*) ^53^ is substantially increased. Values are means ± S.D. from 3 independent experiments. Midguts analyzed 4 days post-induction. **b-d,** The number of *upd3-lacZ*^+^ enterocytes in the R4ab compartment is unchanged by enterocyte *E-cad* depletion (unpaired t-test: *p*>0.05). **e-g,** The number of *10XSTAT-GFP*^*+*^ diploid cells in R4ab is unchanged by enterocyte *E-cad* depletion (unpaired t-test: *p*>0.05). Occasional activation of *10XSTAT-GFP*^*+*^ occurs in *E-cad*-depleted enterocytes (asterisk in **f**; unpaired t-test *p*=0.003), consistent with elevated *Socs36E* (panel **a**). **h-j,** The number of *cycE*^+^ diploid cells in R4ab increases by 92% following enterocyte *E-cad* depletion (unpaired t-test: *p*=0.005). In panels **d**, **g**, and **j**, values are means ± S.D of 4 midguts, analyzed 2 days post-induction. **k,** Expression of *upd3* is not associated with physiological apoptosis. Most enterocytes (~91%) that express *upd3-lacZ* are non-apoptotic, as assessed by staining for cleaved Caspase-3. Values are means ± S.D of 4 midguts, analyzed 6 days post-eclosion. Representative images shown in all panels. All scale bars are 25 μm.

**Extended Data Fig. 7.**
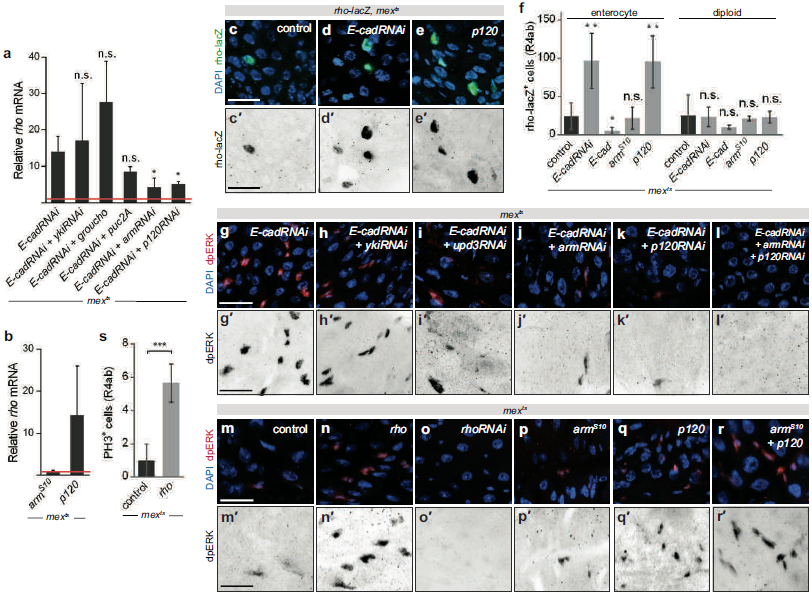
Two E-cad-associated transcription factors, Armadillo and p120-catenin, activate *rho* following loss of *E-cad* in enterocytes. **a,** Enterocyte *armadillo* (*arm*) and *p120-catenin* (*p120*), but not *yorkie* (*yki*) or *groucho*, are necessary for activation of *rho* upon depletion of enterocyte *E-cad*. *rho* mRNAs were measured by qPCR of either *mex ts* control (red line) or *mex ts >E-cadRNAi* midguts, the latter with additional manipulation of candidate *rho* regulators as indicated. Five candidates were examined: Yki, a transcriptional co-activator in the Hpo pathway; Groucho, a co-repressor known to target *rho* in some tissues; JNK, which can augment EGF signaling; and Arm and p120, co-activators that are inhibited by sequestration at E-cad adherens junctions. Neither *yki* depletion nor *groucho* over-expression prevents *rho* activation in *E-cad* knockdown midguts. Overexpression of the JNK in-hibitor *puckered* (*puc2A*) partially reduces *rho* activation, although with unclear significance (unpaired t-test, *p*=0.22). By contrast, knockdown of either *arm* or *p120* significantly reduces *rho* activation (*p*=0.04 and 0.03, respectively). **b,** Overexpression of *p120,* but not *arm*^*S10*^, in enterocytes is sufficient to increase *rho* mRNAs relative to control (red line), as measured by qPCR. In **a** and **b**, values are means ± S.D. of 3 independent experiments; midguts analyzed 4 days post-induction. **c-e**, Depletion of *E-cad* or overexpression of *p120* induces *rho-lacZ* in enterocytes. **f,** Quantification of *rho-lacZ*^+^ cells. The number of *rho-lacZ*^+^ enterocytes increases with *E-cad* de-pletion or *p120* overxpression (unpaired t-test, *p*=0.003 and 0.007 respectively), decreases with *E-cad* overexpression (*p*=0.04), and is unchanged by overexpression of *arm*^*S10*^. The number of *rho-lacZ*^+^ diploid cells is unchanged. In **c-f**, values are means ± S.D of 4 midguts, analyzed 2 days post-induction. **g-l,** Enterocyte *arm* and *p120*, but not *yki* or *upd3*, are necessary for activation of stem cell EGFR following loss of *E-cad*. **g-i,** dpERK^+^ cells are similarly abundant upon double enterocyte RNAi of *E-cad* and either *yki* or *upd3* as upon single RNAi of *E-cad* alone. **j-l,** By contrast, dpERK^+^ cells are substantially decreased upon double RNAi of *E-cad* and either *arm* or *p120* and virtually disappear upon triple RNAi of *E-cad, arm*, and *p120*. **m-o,** Enterocyte *rho* is necessary and sufficient for activation of stem cell EGFR. Overexpression of *rho* in enterocytes increases the abundance of dpERK^+^ stem cells relative to control, whereas depletion of *rho* nearly eliminates them. **p-r,** Enterocyte *p120*, but not *arm*, is sufficient to activate stem cell EGFR. Overexpression of *p120*, but not *arm*^*S10*^, increases the abundance of dpERK^+^ stem cells compared to control. Overexpression of both *p120* and *arm*^*S10*^ together resembles *p120* alone. Panels **g-r** represent two independent experiments; N=4 midguts per genotype, analyzed 2 days after transgene expression. See also Extended Data Fig. 3b. **s,** Overexpression of enterocyte *rho* increases the number of mitotic (phospho-histone H3^+^) stem cells (unpaired t-test, *p*=0.0009). N=4 midguts, assessed after 2 days of transgene expression. Representative images shown in all panels. All scale bars are 25 μm.

**Extended Data Fig. 8.**
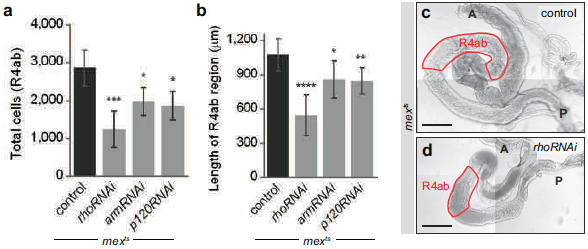
Loss of *rho, arm*, or *p120* in enterocytes results in organ atrophy. **a,** Total R4ab cell counts. Depletion of *rho* in enterocytes reduces total cells by 60% compared to control (unpaired t-test, *p*=0.0007). Depletion of either *arm* or *p120* reduces total cells by ~35% (*p*=0.011 and *p*=0.012, respectively). Values are means ± S.D from one of three representative experiments. N=4 midguts per genotype, analyzed after 6 days of induction. **b**-**d,** Depletion of enterocyte *rho, arm*, or *p120* reduces organ size. The R4ab compartment is significantly shorter following depletion of enterocyte *rho, arm*, or *p120* compared to control (unpaired t-test, *p*<0.0001, 0.011, and 0.0001 respectively). N=10-12 midguts per genotype, analyzed after 6 days of induction. Representative images are shown. A, anterior; P posterior. Scale bars: 200μm.

## REFERENCES

1. Leblond, C. P. & Stevens, C. E. The constant renewal of the intestinal epithelium in the albino rat. Anat. Rec. 100, 357–377 (1948).

2. Pellettieri, J. & Alvarado, A. S. Cell turnover and adult tissue homeostasis: From humans to planarians. Annu Rev Genet 41, 83–105 (2007).

3. O’Brien, L.E. & Bilder, D. Beyond the niche: Tissue-level coordination of stem cell dynamics. Annu. Rev. Cell. Dev. Biol. 29, 107–136 (2013).

4. Siudeja, K. et al. Frequent somatic mutation in adult intestinal stem cells drives neoplasia and genetic mosaicism during aging. Cell Stem Cell 17, 663–674 (2015).

5. Alcolea, M. P. et al. Differentiation imbalance in single oesophageal progenitor cells causes clonal immortalization and field change. Nature Cell Biology 16, 615–622 (2014).

6. O’Brien, L. E., Soliman, S. S., Li, X. & Bilder, D. Altered modes of stem cell division drive adaptive intestinal growth. Cell 147, 603–614 (2011).

7. Visvader, J. E. Cells of origin in cancer. Nature 469, 314–322 (2011).

8. Micchelli, C. A. & Perrimon, N. Evidence that stem cells reside in the adult Drosophila midgut epithelium. Nature 439, 475–479 (2006).

9. Ohlstein, B. & Spradling, A. The adult Drosophila posterior midgut is maintained by pluripotent stem cells. Nature 439, 470–474 (2006).

10. Clermont, Y. & Leblond, C. P. Renewal of spermatogonia in the rat. Am. J. Anat. 93, 475–501 (1953).

11. Biteau, B., Hochmuth, C. E. & Jasper, H. Maintaining tissue homeostasis: dynamic control of somatic stem cell activity. Cell Stem Cell 9, 402–411 (2011).

12. Clevers, H. What is an adult stem cell? Science 350, 1319–1320 (2015).

13. Jiang, H. et al. Cytokine/Jak/Stat signaling mediates regeneration and homeostasis in the Drosophila midgut. Cell 137, 1343–1355 (2009).

14. Buchon, N. et al. Morphological and molecular characterization of adult midgut compartmentalization in Drosophila. Cell Rep 3, 1725–1738 (2013).

15. Marianes, A. & Spradling, A. C. Physiological and stem cell compartmentalization within the Drosophila midgut. eLife 2, e00886 (2013).

16. Ohlstein, B. & Spradling, A. Multipotent Drosophila intestinal stem cells specify daughter cell fates by differential notch signaling. Science 315, 988–992 (2007).

17. Hudry, B., Khadayate, S. & Miguel-Aliaga, I. The sexual identity of adult intestinal stem cells controls organ size and plasticity. Nature 530, 344–348 (2016).

18. Chen, H., Zheng, X. & Zheng, Y. Age-associated loss of lamin-B leads to systemic inflammation and gut hyperplasia. Cell 159, 829–843 (2014).

19. Harrison, D. A. & Perrimon, N. Simple and efficient generation of marked clones in Drosophila. 3, 424–433 (1993).

20. de Navascués, J. et al. Drosophila midgut homeostasis involves neutral competition between symmetrically dividing intestinal stem cells. EMBO J 31, 2473–2485 (2012).

21. Ayyaz, A., Li, H. & Jasper, H. Haemocytes control stem cell activity in the Drosophila intestine. Nature Cell Biology 17, 736–748 (2015).

22. Takeishi, A. et al. Homeostatic epithelial renewal in the gut is required for dampening a fatal systemic wound response in Drosophila. Cell Rep 3, 919–930 (2013).

23. Kessler, T. & Müller, H. A. J. Cleavage of Armadillo/beta-catenin by the caspase DrICE in Drosophila apoptotic epithelial cells. BMC Dev. Biol. 9, 15 (2009).

24. Steinhusen, U. et al. Cleavage and shedding of E-cadherin after induction of apoptosis. J. Biol. Chem. 276, 4972–4980 (2001).

25. Keller, S. H. & Nigam, S. K. Biochemical processing of E-cadherin under cellular stress. Biochemical and Biophysical Research Communications 307, 215–223 (2003).

26. Hung, C.-F., Chiang, H.-S., Lo, H.-M., Jian, J.-S. & Wu, W.-B. E-cadherin and its downstream catenins are proteolytically cleaved in human HaCaT keratinocytes exposed to UVB. Exp. Dermatol. 15, 315–321 (2006).

27. Jeanes, A., Gottardi, C. J. & Yap, A. S. Cadherins and cancer: how does cadherin dysfunction promote tumor progression? Oncogene 27, 6920–6929 (2008).

28. Hermiston, M. L. & Gordon, J. I. In vivo analysis of cadherin function in the mouse intestinal epithelium: Essential roles in adhesion, maintenance of differentiation, and regulation of programmed cell death. The Journal of Cell Biology 129, 489–506 (1995).

29. Watabe, M., Nagafuchi, A., Tsukita, S. & Takeichi, M. Induction of polarized cell-cell association and retardation of growth by activation of the E-cadherin-catenin adhesion system in a dispersed carcinoma line. The Journal of Cell Biology 127, 247–256 (1994).

30. Kim, N.-G., Koh, E., Chen, X. & Gumbiner, B. M. E-cadherin mediates contact inhibition of proliferation through Hippo signaling-pathway components. Proceedings of the National Academy of Sciences 108, 11930–11935 (2011).

31. McClatchey, A. I. & Yap, A. S. Contact inhibition (of proliferation) redux. Curr Opin Cell Biol 24, 685–694 (2012).

32. Huang, J., Zhou, W., Dong, W., Watson, A. M. & Hong, Y. Directed, efficient, and versatile modifications of the Drosophila genome by genomic engineering. Proceedings of the National Academy of Sciences 106, 8284–8289 (2009).

33. Kolahgar, G. et al. Cell Competition Modifies Adult Stem Cell and Tissue Population Dynamics in a JAK-STAT-Dependent Manner. Developmental Cell 34, 297–309 (2015).

34. Campbell, K. & Casanova, J. A role for E-cadherin in ensuring cohesive migration of a heterogeneous population of non-epithelial cells. Nat Commun 6, 7998 (2015).

35. Maeda, K., Takemura, M., Umemori, M. & Adachi-Yamada, T. E-cadherin prolongs the moment for interaction between intestinal stem cell and its progenitor cell to ensure Notch signaling in adult Drosophila midgut. Genes to Cells 13, 1219–1227 (2008).

36. Choi, N.-H., Lucchetta, E. & Ohlstein, B. Nonautonomous regulation of Drosophila midgut stem cell proliferation by the insulin-signaling pathway. Proceedings of the National Academy of Sciences 108, 18702–18707 (2011).

37. Nelson, W. J. & Nusse, R. Convergence of Wnt, beta-catenin, and cadherin pathways. Science 303, 1483–1487 (2004).

38. Yang, C.-C. et al. Differential regulation of the Hippo pathway by adherens junctions and apical-basal cell polarity modules. Proceedings of the National Academy of Sciences 112, 1785–1790 (2015).

39. Geletu, M., Guy, S., Arulanandam, R., Feracci, H. & Raptis, L. Engaged for survival: From cadherin ligation to STAT3 activation. JAKSTAT 2, e27363 (2013).

40. Dumstrei, K., Wang, F., Shy, D., Tepass, U. & Hartenstein, V. Interaction between EGFR signaling and DE-cadherin during nervous system morphogenesis. Development 129, 3983–3994 (2002).

41. Lu, L. et al. LRIG1 regulates cadherin-dependent contact inhibition directing epithelial homeostasis and pre-invasive squamous cell carcinoma development. J Pathol 229, 608–620 (2013).

42. Takahashi, K. & Suzuki, K. Density-dependent inhibition of growth involves prevention of EGF receptor activation by E-cadherin-mediated cell-cell adhesion. Exp Cell Res 226, 214–222 (1996).

43. Takashima, S. & Hartenstein, V. Genetic control of intestinal stem cell specification and development: a comparative view. Stem Cell Rev 8, 597–608 (2012).

44. Biteau, B. & Jasper, H. EGF signaling regulates the proliferation of intestinal stem cells in Drosophila. Development 138, 1045–1055 (2011).

45. Lucchetta, E. M. & Ohlstein, B. The Drosophila midgut: a model for stem cell driven tissue regeneration. WIREs Dev Biol 1, 781–788 (2012).

46. Jiang, H., Grenley, M. O., Bravo, M.-J., Blumhagen, R. Z. & Edgar, B. A. EGFR/Ras/MAPK signaling mediates adult midgut epithelial homeostasis and regeneration in Drosophila. Cell Stem Cell 8, 84–95 (2011).

47. Cordero, J. B., Stefanatos, R. K., Myant, K., Vidal, M. & Sansom, O. J. Non-autonomous crosstalk between the Jak/Stat and Egfr pathways mediates Apc1-driven intestinal stem cell hyperplasia in the Drosophila adult midgut. Development 139, 4524–4535 (2012).

48. Zhou, F., Rasmussen, A., Lee, S. & Agaisse, H. The UPD3 cytokine couples environmental challenge and intestinal stem cell division through modulation of JAK/STAT signaling in the stem cell microenvironment. Developmental Biology 373, 383–393 (2013).

49. Osman, D. et al. Autocrine and paracrine unpaired signaling regulate intestinal stem cell maintenance and division. Journal of Cell Science 125, 5944–5949 (2012).

50. Jiang, H. & Edgar, B. A. EGFR signaling regulates the proliferation of Drosophila adult midgut progenitors. Development 136, 483–493 (2009).

51. Xu, N. et al. EGFR, Wingless and JAK/STAT signaling cooperatively maintain Drosophila intestinal stem cells. Developmental Biology 354, 31–43 (2011).

52. Buchon, N., Broderick, N. A., Kuraishi, T. & Lemaitre, B. Drosophila EGFR pathway coordinates stem cell proliferation and gut remodeling following infection. BMC Biol 8, 152 (2010).

53. Jin, Y. et al. EGFR/Ras signaling controls Drosophila intestinal stem cell proliferation via Capicua-regulated genes. PLoS Genetics 11, e1005634 (2015).

54. Strand, M. & Micchelli, C. A. Regional control of Drosophila gut stem cell proliferation: EGF establishes GSSC proliferative set point & controls emergence from quiescence. PLoS ONE 8, e80608 (2013).

55. Qian, X., Karpova, T., Sheppard, A. M., McNally, J. & Lowy, D. R. E-cadherin-mediated adhesion inhibits ligand-dependent activation of diverse receptor tyrosine kinases. EMBO J 23, 1739–1748 (2004).

56. Guo, Z. et al. E-cadherin interactome complexity and robustness resolved by quantitative proteomics. Sci Signal 7, rs7–rs7 (2014).

57. Roy, S., Hsiung, F. & Kornberg, T. B. Specificity of Drosophila cytonemes for distinct signaling pathways. Science 332, 354–358 (2011).

58. Lander, A. D. Morpheus unbound: reimagining the morphogen gradient. Cell 128, 245–256 (2007).

59. Lee, T. & Luo, L. Mosaic analysis with a repressible cell marker for studies of gene function in neuronal morphogenesis. Neuron 22, 451–461 (1999).

60. Lee, J. R., Urban, S., Garvey, C. F. & Freeman, M. Regulated intracellular ligand transport and proteolysis control EGF signal activation in Drosophila. Cell 107, 161–171 (2001).

61. Urban, S., Lee, J. R. & Freeman, M. A family of Rhomboid intramembrane proteases activates all Drosophila membrane-tethered EGF ligands. EMBO J 21, 4277–4286 (2002).

62. McCrea, P. D., Gu, D. & Balda, M. S. Junctional music that the nucleus hears: cell-cell contact signaling and the modulation of gene activity. Cold Spring Harb Perspect Biol 1, a002923–a002923 (2009).

63. Du, W. et al. From cell membrane to the nucleus: An emerging role of E-cadherin in gene transcriptional regulation. J. Cell. Mol. Med. 18, 1712–1719 (2014).

64. Daniel, J. M. & Reynolds, A. B. The catenin p120(ctn) interacts with Kaiso, a novel BTB/POZ domain zinc finger transcription factor. Mol Cell Biol 19, 3614–3623 (1999).

65. Kelly, K. F. NLS-dependent nuclear localization of p120ctn is necessary to relieve Kaisomediated transcriptional repression. Journal of Cell Science 117, 2675–2686 (2004).

66. Park, J.-I. et al. Kaiso/p120-catenin and TCF/beta-catenin complexes coordinately regulate canonical Wnt gene targets. Developmental Cell 8, 843–854 (2005).

67. Hosking, C. R. et al. The Transcriptional Repressor Glis2 Is a Novel Binding Partner for p120 Catenin. Mol Biol Cell 18, 1918–1927 (2007).

68. Lee, M., Ji, H., Furuta, Y., Park, J.-I. & McCrea, P. D. p120-catenin regulates REST and CoREST, and modulates mouse embryonic stem cell differentiation. Journal of Cell Science 127, 4037–4051 (2014).

69. Cavallo, R. A. et al. Drosophila Tcf and Groucho interact to repress Wingless signalling activity. Nature 395, 604–608 (1998).

70. Daniels, D. L. & Weis, W. I. Beta-catenin directly displaces Groucho/TLE repressors from Tcf/Lef in Wnt-mediated transcription activation. Nat. Struct. Mol. Biol. 12, 364–371 (2005).

71. Zhang, T. & Du, W. Groucho restricts rhomboid expression and couples EGFR activation with R8 selection during Drosophila photoreceptor differentiation. Developmental Biology 407, 246–255 (2015).

72. Desai, T. J., Brownfield, D. G. & Krasnow, M. A. Alveolar progenitor and stem cells in lung development, renewal and cancer. 507, 190–194 (2014).

73. Lei, K. et al. Egf signaling directs neoblast repopulation by regulating asymmetric cell division in planarians. Developmental Cell 38, 413–429 (2016).

74. Gilboa, L. & Lehmann, R. Soma-germline interactions coordinate homeostasis and growth in the Drosophila gonad. Nature 443, 97–100 (2006).

75. Abba, M. C. et al. Rhomboid domain containing 2 (RHBDD2): a novel cancer-related gene over-expressed in breast cancer. Biochim Biophys Acta 1792, 988–997 (2009).

76. Yan, Z. et al. Human rhomboid family-1 gene silencing causes apoptosis or autophagy to epithelial cancer cells and inhibits xenograft tumor growth. Mol. Cancer Ther. 7, 1355–1364 (2008).

77. Zou, H. et al. Human rhomboid family-1 gene RHBDF1 participates in GPCR-mediated transactivation of EGFR growth signals in head and neck squamous cancer cells. FASEB J. 23, 425–432 (2009).

78. Song, W. et al. Rhomboid domain containing 1 promotes colorectal cancer growth through activation of the EGFR signalling pathway. Nat Commun 6, 8022 (2015).

## Extended Data References (not cited in main text)

79. Oda, H. & Tsukita, S. Nonchordate classic cadherins have a structurally and functionally unique domain that is absent from chordate classic cadherins. Developmental Biology 216, 406–422 (1999).

80. Freeman, M., Kimmel, B. E. & Rubin, G. M. Identifying targets of the rough homeobox gene of Drosophila: evidence that rhomboid functions in eye development. Development 116, 335–346 (1992).

81. O’Brien, L. E. Regional specificity in the Drosophila midgut: setting boundaries with stem cells. Cell Stem Cell 13, 375–376 (2013).

82. Strand, M. & Micchelli, C. A. Quiescent gastric stem cells maintain the adult Drosophila stomach. Proceedings of the National Academy of Sciences 108, 17696–17701 (2011).

83. Cordero, J. B., Stefanatos, R. K., Scopelliti, A., Vidal, M. & Sansom, O. J. Inducible progenitor-derived Wingless regulates adult midgut regeneration in Drosophila. EMBO J 31, 3901–3917 (2012).

84. Zeng, X. & Hou, S. X. Enteroendocrine cells are generated from stem cells through a distinct progenitor in the adult Drosophila posterior midgut. Development 142, 644–653 (2015).

85. Antonello, Z. A., Reiff, T., Ballesta-Illan, E. & Dominguez, M. Robust intestinal homeostasis relies on cellular plasticity in enteroblasts mediated by miR-8-Escargot switch. EMBO J 34, 2025–2041 (2015).

86. Takashima, S., Younossi-Hartenstein, A., Ortiz, P. A. & Hartenstein, V. A novel tissue in an established model system: the Drosophila pupal midgut. Dev Genes Evol 221, 69–81 (2011).

87. Rera, M. et al. Modulation of longevity and tissue homeostasis by the Drosophila PGC-1 homolog. Cell Metab 14, 623–634 (2011).

88. Shaw, R. L. et al. The Hippo pathway regulates intestinal stem cell proliferation during Drosophila adult midgut regeneration. Development 137, 4147–4158 (2010).

89. Biteau, B. & Jasper, H. Slit/Robo signaling regulates cell fate decisions in the intestinal stem cell lineage of Drosophila. Cell Rep 7, 1867–1875 (2014).

90. Eroglu, M. & Derry, W. B. Your neighbours matter –non-autonomous control of apoptosis in development and disease. Cell Death Differ 23, 1110–1118 (2016).

91. Tian, A., Benchabane, H., Wang, Z. & Ahmed, Y. Regulation of stem cell proliferation and cell fate specification by Wingless/Wnt signaling gradients enriched at adult intestinal compartment boundaries. PLoS Genetics 12, e1005822 (2016).

92. DasGupta, R., Kaykas, A., Moon, R. T. & Perrimon, N. Functional genomic analysis of the Wnt-wingless signaling pathway. Science 308, 826–833 (2005).

93. Karpowicz, P., Perez, J. & Perrimon, N. The Hippo tumor suppressor pathway regulates intestinal stem cell regeneration. Development 137, 4135–4145 (2010).

94. Ren, W. et al. Windpipe controls Drosophila intestinal homeostasis by regulating JAK/STAT pathway via promoting receptor endocytosis and lysosomal degradation. PLoS Genetics 11, e1005180 (2015).

